# Allochthony in stream food webs decreases with temperature across a subcontinental scale

**DOI:** 10.64898/2026.09.27.752912

**Authors:** Kierstyn T Higgins, Michael T. Bogan, Albert Ruhí, Meryl C. Mims, Carla L. Atkinson, Michelle H. Busch, Brian A. Gill, Chelsea R. Smith, Thomas M. Neeson, Yang Hong, Travis M. Apgar, Arial J. Shogren, Megan C. Malish, Samuel Silknetter, Rose M. Mohammadi, Alice M. Belskis, Daniel C. Allen

**Author notes:** **Corresponding Author:** Kierstyn T. Higgins, 236 Forest Resources Building, 466 Bigler Rd., The Pennsylvania State University, University Park, PA, USA 16802. **Statement of authorship:** DCA, AR, TMN, AJS, CLA, MTB, MCM and YH: conceptualization. KTH, BAG, MTB, TMA, AMB, and DCA: data curation. KTH, MHB, MCM, RMM, BAG, CRS, AJS, CLA, MTB, TMA, AR, SS, MM, AMB, YH and DCA: investigation. KTH and DCA: methodology. TMN, CLA, MTB, AR, MCM and DCA: project administration. KTH and MHB: software. KTH and DCA: resources. KTH, TMN, CLA, MM, MTB, AR, MCM and DCA: supervision. KTH: validation and visualization. KTH and DCA: formal data analysis. KTH: original draft of the manuscript. All authors contributed to subsequent drafts of the manuscript. TMN and DCA: funding acquisition. **Data accessibility statement:** Data and code are available on GitHub (https://github.com/KierstynHiggins/manuscript-temperature-allochthony/tree/main).

## Abstract

Riverine and riparian food webs are connected via “allochthonous” resource flows from neighboring ecosystems. These resources, such as terrestrial leaf litter, contribute to benthic macroinvertebrate diets, supporting higher trophic levels. While allochthony occurs in streams globally, its sensitivity to broad environmental gradients remains poorly understood. We explored the drivers of allochthony across six river basins within the contiguous United States, including climate, stream size, and flow permanence. We used Bayesian mixing models to estimate macroinvertebrate dietary proportions in four functional feeding groups. Air temperature best predicted allochthony, with consumers relying less on allochthonous resources with increasing temperature. Our models indicate that macroinvertebrate diets become primarily autochthonous at approximately 15°C mean annual air temperature, with leaf litter-consuming shredders showing a slightly higher threshold for this shift. We also found increased allochthony in smaller, non-perennial streams. These findings highlight the complexity of stream allochthony and responses to multi-scale environmental factors.

## INTRODUCTION

Since the late twentieth century, ecological theory has recognized streams as open, non-equilibrial ecosystems (Junk *et al*. 1989; Minshall *et al*. 1985; Vannote *et al*. 1980) whose food webs are structured by the dynamic transfer of materials to and from neighboring habitats (Baxter *et al*. 2004; Forrester *et al*. 1999; Lindeman 1942). This exchange of energy and materials results in organisms within the stream assimilating resources that originate from another ecosystem (and vice versa), a phenomenon known as “allochthony” (Nakano & Murakami 2001; Webster & Benfield 1986). The relative importance of terrestrial leaf litter as a food source in stream food webs, particularly in temperate headwaters, is well established (Anderson & Sedell 1979; Graça 2001; Vannote *et al*. 1980). The extent to which allochthony in streams varies across climates remains unresolved, and existing research has yielded contrasting results (Allen *et al*. 2024). Therefore, understanding the drivers of allochthony in streams is still a key goal in modern ecology.

To date, the most widely recognized driver of allochthony is stream size. As stream size increases, canopy cover decreases, light penetration increases and supports greater *in situ* primary production (“autochthonous” energy), while terrestrially derived (“allochthonous”) energy inputs decrease (Collins *et al*. 2016; Minshall 1978; Vannote *et al*. 1980). This foundational concept forms The River Continuum Concept (RCC; Vannote *et al*. 1980), yet subsequent theories posit other influences on allochthony (Junk *et al*. 1989; Montgomery 1999; Thorp & Delong 1994). The Freshwater Biome Gradient Framework, for example, highlights broad-scale patterns in streams across latitudinal and elevational gradients, proposing that climate controls stream food webs and allochthony (Dodds *et al*. 2015, 2019). Climatic variation in temperature, precipitation, and hydrologic disturbances should strongly affect streams, such that those in cool and wet biomes should rely more on allochthonous inputs while streams in warm and dry biomes should rely more on autochthonous carbon (Dodds *et al*. 2015). Streams in cool and wet biomes are generally more shaded from the riparian overstory and have greater allochthonous production, while warm and dry biomes have shorter and more sparse riparian vegetation, resulting in more sunlight and higher autochthonous production. In addition, the Freshwater Biome Gradient Framework (Dodds *et al*. 2019) also highlights the climate-related hydrologic disturbances that can result in local-scale stream drying.

Streams that experience periodic flow cessation (hereafter referred to as “non-perennial streams”) approach 70 percent of global stream length (Botter *et al*. 2026), and can range from long stretches of non-perennial streams and rivers in arid climates (Caruso & Haynes 2011) to localized intermittent headwaters in temperate climates (Lake 2003). Non-perennial streams harbor unique, desiccation-resistant taxa (Larned *et al*. 2010), as well as strong dispersers that can rapidly recolonize wetted habitats (Sarremejane *et al*. 2017). Non-perennial streams also accumulate large amounts of terrestrial detritus which is later processed or exported upon the rewetting of stream channels, contributing to downstream metabolism and production (Datry *et al*. 2018; Steward *et al*. 2012). Despite their prevalence, current conceptual models understate the importance of drying in stream ecosystem dynamics, including allochthony (Allen *et al*. 2020).

Beyond resource availability, consumer resource preferences are important for determining reliance on allochthonous inputs. Assimilation of resources by aquatic invertebrates is often a function of metabolic needs that differ depending on functional traits and external pressures such as thermal stress (Spooner & Vaughn 2008). Additionally, available food sources may differ in nutritional quality. Allochthonous sources are generally high in carbon and low in nitrogen (high C:N ratio), while autochthonous sources require less structural carbon, resulting in a lower C:N ratio (Evans-White & Halvorson 2017; Lau *et al*. 2009). C:N ratios of sources are used as an indicator of food quality for macroinvertebrates, as a lower C:N ratio reflects higher nitrogen (protein) content relative to carbon and is generally preferred by consumers (Frost *et al*. 2002). Variation in resource stoichiometry can also be driven by environmental factors. For example, the same plant species can have significantly differing C:N ratios depending on factors such as light availability and water availability (Cronin & Lodge 2003). Therefore, allochthony in streams is a two-fold process that intertwines resource availability and organismal traits.

Aquatic macroinvertebrates are the dominant primary consumers in streams and serve as an intermediate link between higher trophic levels and basal resources (Wallace & Webster 1996). Macroinvertebrates fulfill different functional roles according to their feeding traits, also known as functional feeding groups (FFG; Wallace & Webster 1996). Important FFGs include collector-gatherers that consume deposited organic matter; collector-filterers that collect suspended algal cells or detritus; herbivores that scrape algae or graze on macrophytes; and shredders that break down coarse particulate organic matter such as leaf litter (Twardochleb *et al*. 2021). Thus, different FFGs should differ in their reliance on allocthonous vs. autochothonus materials; herbivores should have more autochthonous contributions to their diet while shredders should have more allochthonous contributions.

Determining relative proportions of allochthonous and autochthonous resources in consumer diets is challenging when direct observation of feeding behavior is not feasible. Stable isotope analysis overcomes this challenge. Carbon (C) and Nitrogen (N) stable isotopes can be used to study allochthony in stream food webs (Layman *et al*. 2012), as δ^15^N values increase with trophic level (Minagawa & Wada 1984; Peterson & Fry 1987), while δ^13^C values vary based on photosynthetic pathways (DeNiro & Epstein 1981). Freshwater primary producers are typically enriched in δ^13^C relative to terrestrial plants (Milligan *et al*. 2010). In addition to trophic discrimination, δ^15^N can also assist in discriminating between sources, as different sources may have different N-uptake mechanisms (Mulholland *et al*. 2000).

Using stable isotope analysis, we estimated macroinvertebrate dietary proportions of allochthonous to autochthonous resources across a broad spatial and climatic scale, with the goal of detecting broad-scale patterns of allochthony across a longitudinal gradient. We hypothesized that 1) as stream size increases, allochthonous contributions to consumer diets should decrease due to reduced canopy cover and terrestrial inputs, 2) allochthony should be less important in streams in arid climates due to a lack of canopy cover and increased light availability, 3) streams with increased drying within a watershed would rely more on allochthonous carbon, due to the decrease in autochthonous primary production as a result of stream drying, 4) increased allochthonous source quality (lower C:N ratio) will lead to increased allochthony, and 5) dietary proportions will be distinct across FFGs.

## MATERIALS AND METHODS

### Study Sites

We chose 6 stream networks representing the southern US climatic gradient (Figure 1). Within each stream network, we selected 6–10 study sites that capture the range of diversity of streams in the network (channel morphology, flow regime, riparian zones, etc.) and including a National Ecological Observatory Network (NEON; Metzger *et al*. 2019) site when possible, for a total of 50 sites (Table 1). The Kings Creek, KS (KING), Blue River, OK (BLUE), and West Prong Little Pigeon, TN (WPLP) basins were sampled in Spring 2021 and the Sycamore Creek, AZ (SYCA), Mayfield Creek, AL (MAYF), and Chalone Creek, CA (PINN) basins were sampled in Spring 2022. We elected to sample in the spring, typically the wettest time of year when non-perennial reaches contained flowing water.

**Figure 1.**
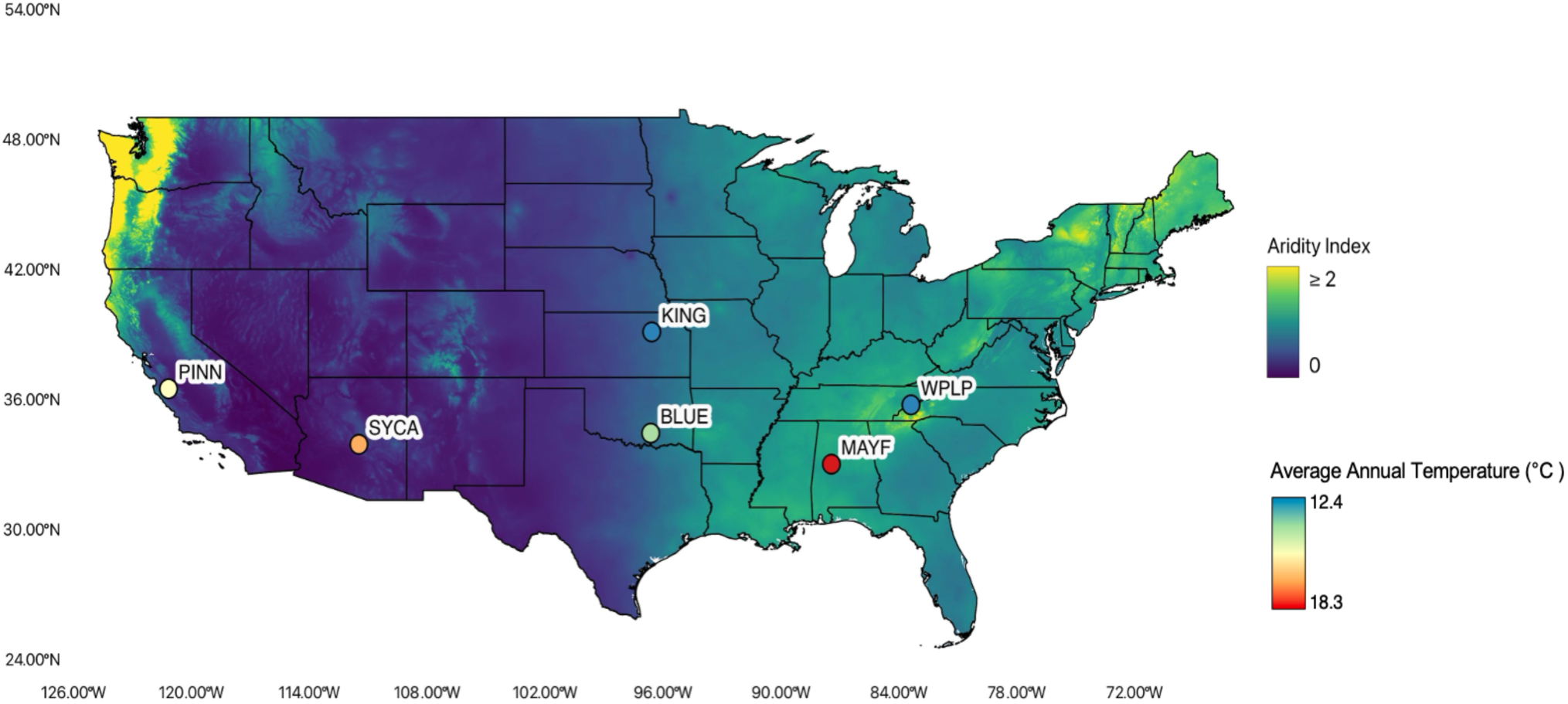
Aridity index map of the contiguous US with state borders (Zomer *et al*. 2022). The six study locations are colored by average annual temperature gathered from the PRISM Group at Oregon State University (“PRISM Group at Oregon State University” 2026).

**Table 1.** Table of all environmental variables measured for each basin, including the number of sites, mean **±** standard deviation of all sites within a basin and the range of all sites within a basin.

| Basin | Name | (n) Site | Precipitation (mm) | Temperature (°C) | Aridity Index | Proportion of Dry Days | % Canopy Cover | Wetted Width (m) | Drainage Area (km<sup>2</sup>) | CN Ratio |
| --- | --- | --- | --- | --- | --- | --- | --- | --- | --- | --- |
| **BLUE** | Blue River, OK, USA | 9 | 1014.57 ± 14.49<br>(991.69–1037.51) | 15.62 ± 0.05<br>(15.56–15.68) | 0.71 ± 0.00<br>(0.70–0.71) | 0.41 ± 0.20<br>(0.11–0.68) | 68.87 ± 33.07<br>(0.00–94.80) | 3.99 ± 3.35<br>(1.00–10.94) | 9.68 ± 18.99<br>(0.37–59.49) | 28.07 ± 5.96<br>(19.75–39.65) |
| **KING** | Kings Creek, KS, USA | 10 | 843.33 ± 6.78<br>(832.03–852.32) | 12.39 ± 0.08<br>(12.21–12.47) | 0.66 ± 0.00<br>(0.65–0.66) | 0.37 ± 0.26<br>(0.01–0.71) | 75.34 ± 11.78<br>(59.20–95.80) | 3.38 ± 1.98<br>(1.76–8.10) | 3.39 ± 4.91<br>(0.29–16.62) | 30.41 ± 5.46<br>(21.06–36.83) |
| **MAYF** | Mayfield Creek, AL, USA | 9 | 1401.13 ± 11.31<br>(1383.21–1419.04) | 18.24 ± 0.06<br>(18.14–18.33) | 1.14 ± 0.00<br>(1.13–1.15) | 0.01 ± 0.02<br>(0.00–0.07) | 73.08 ± 11.02<br>(53.00–83.10) | 4.09 ± 3.67<br>(1.47–12.90) | 15.26 ± 27.97<br>(0.12–85.81) | 13.62 ± 1.23<br>(11.62–15.10) |
| **PINN** | Chalone Creek, CA, USA | 6 | 217.78 ± 12.11<br>(203.79–236.05) | 16.15 ± 0.24<br>(15.90–16.54) | 0.27 ± 0.05<br>(0.23–0.32) | 0.29 ± 0.26<br>(0.02–0.58) | 37.40 ± 33.34<br>(2.10–93.80) | 1.93 ± 1.04<br>(0.95–3.75) | 57.40 ± 51.46<br>(11.64–140.22) | 21.60 ± 12.95<br>(7.61–38.97) |
| **SYCA** | Sycamore Creek, AZ, USA | 7 | 487.21 ± 77.17<br>(377.33–601.14) | 17.21 ± 1.98<br>(14.94–20.59) | 0.30 ± 0.00<br>(0.22–0.36) | 0.30 ± 0.23<br>(0.00–0.60) | 47.57 ± 24.30<br>(20.00–90.00) | 3.66 ± 4.62<br>(0.55–13.86) | 57.07 ± 70.85<br>(2.24–191.00) | 18.39 ± 4.61<br>(12.12–24.34) |
| **WPLP** | West Prong Little Pigeon River, TN, USA | 9 | 1808.72 ± 222.22<br>(1487.91–2114.17) | 12.83 ± 0.92<br>(11.58–14.32) | 1.28 ± 0.12<br>(1.15–1.54) | 0.08 ± 0.08<br>(0.00–0.20) | 85.86 ± 9.48<br>(69.00–95.60) | 2.42 ± 1.45<br>(0.96–4.80) | 0.57 ± 0.62<br>(0.09–1.84) | 30.98 ± 4.44<br>(24.63–37.84) |

### Environmental variables

At each study site, we established a 150-m study reach. Stream Temperature Intermittence and Conductivity (STIC) sensors were installed every 30 meters across a range of habitats (riffle, run, and pool; Chapin *et al*. 2014), which recorded temperature and relative conductivity at 30–45-minute intervals. We installed five sensors per site to account for potential sensor loss or malfunction. At some sites (KING-07, KING-08, BLUE-09), floods regularly removed STIC sensors, so we installed three game cameras at each site to take daily images of the stream reaches, which were classified as wet or dry based on manual interpretation of the images. We monitored streamflow at these sites from 2019-2024. To determine wet/dry status of each sensor, the raw sensor data were converted to water presence/absence using the R package *STICr* (Zipper *et al*. 2025) or manual data inspection, with individual threshold cutoffs of relative conductivity for determining wet/dry status, depending on the baseline conductivity for each sensor. For each site and day, we calculated the proportion of active sensors that were dry during the 365 days preceding biological sampling. Days were classified as dry when ≥60% of active sensors were dry. We calculated the proportion of dry days for each site using the 365-day sampling period and, to account for temporal gaps and one site lacking data (WPLP10), using all available observations from 2019–2024. A paired t-test detected no difference between estimates (*t*_51_ = -0.66, *p* = 0.51), so we used the full 2019–2024 dataset for analysis.

At 30-meter intervals, we recorded canopy cover (%) using a spherical densiometer and wetted width (m) of the stream channel using a measuring tape, which is a measurement of stream size (Doi 2009; Jonsson *et al*. 2018). SYCA did not have canopy cover or wetted width data for the year sampled, so we used data collected from the year prior except for sites SYCA03, SYCA09, and SYCA10 that did not have wetted width data, so these sites were removed from the analysis. Watersheds were delineated using a drainage threshold of 0.1 km ^2^ and drainage area (km^2^) was calculated using digital elevation models for each site. We downloaded 800 m resolution daily precipitation (mm) and daily temperature (°C) data for each site from The Prism Group database (“PRISM Group at Oregon State University” 2026) for the year prior to the date sampled. We then calculated total annual precipitation and mean annual temperature for each site. Additionally, we obtained annual average aridity index data for each site from the 1970-2000 period, quantified as the ratio of precipitation to potential evapotranspiration (Zomer *et al*. 2022). Correlation coefficients were calculated for all observed environmental variables to assess collinearity (Figure 2). For all environmental variables, we calculated the mean ± standard deviation and range for each basin (Table 1). Further description of basin characteristics can be found in Supplementary Methods: *Summary of environmental variables*.

**Figure 2.**
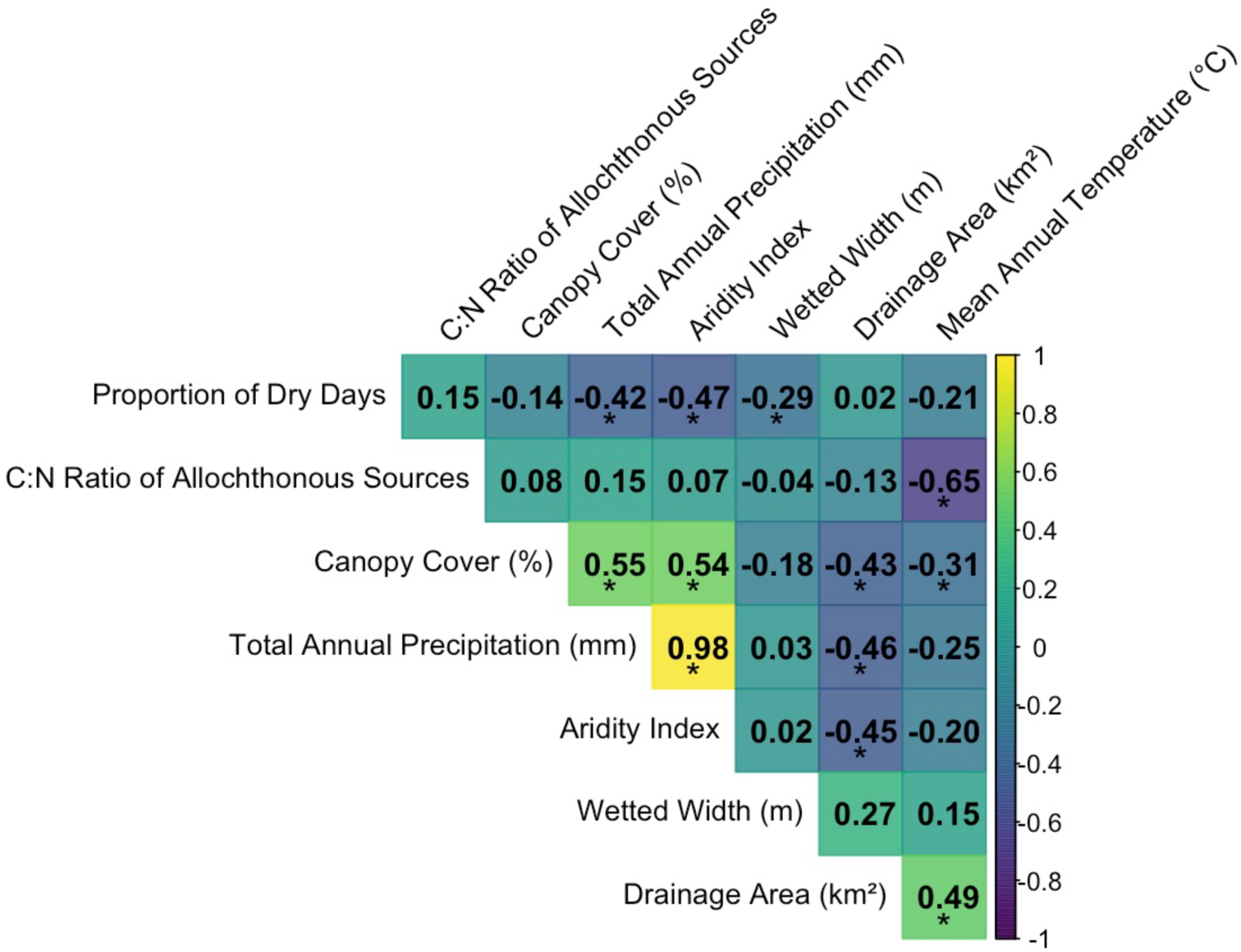
Correlation matrix plot of environmental variables, showing the correlation coefficient in each cell, and colored by correlation coefficient. Cells with (*) indicate a p-value of ≤ 0.05.

### Sampling of basal resources and invertebrates

Leaf litter, filamentous algae, bryophytes and macrophytes were hand collected from each site. Epilithon (biofilm) was sampled from 5 large rocks at each site using a bristle brush to scrape material off the rocks into a slurry. The epilithon slurry was rinsed and put through a 500-micron sieve to remove unwanted material. To collect Fine Benthic Organic Matter (FBOM), an 8–10-inch diameter piece of PVC pipe was placed in the substrate, with the top of the pipe exposed above the water line. Next, we agitated substrate by hand and suspended in the water within the pipe. This suspended material was extracted with a large, bulb-operated syringe. All basal resource samples were placed in whirl-paks, kept on ice and stored at -20℃ when returning to the lab. Invertebrate samples were qualitatively collected using a D-frame net from all potential macrohabitats within a site (i.e. riffles, runs, and pools), sampling for 15 minutes. We sieved and removed large debris from samples in the field and kept them on ice prior to storing at -20℃ when returning to the lab.

### Sample preparation of basal resources for stable isotope analysis

Leaf litter, filamentous algae, and macrophyte samples were lyophilized for 48-72 hours or until completely dry, and ground with a Wiley Mill using a 40-mesh screen. Epilithon and FBOM samples were vacuum filtered through a pre-ashed and pre-weighed 25mm glass microfiber filter (GF/F, Whatman). Filters were oven-dried at 50℃ for 48 hours and weighed. For samples with enough material the contents were scraped off the filter and placed in a vial. For the small number of samples that did not have enough material, a known fraction of the filter was removed using a template, and the weight of the filter piece was subtracted, giving an estimated weight of the sample used, and those samples were processed with the filter portion still intact. All basal resource samples were weighed and packed into 9 x 5 mm or 12 x 6 mm tin capsules (Elemental Microanalysis, Part #: D1022 and D1012, respectively). Target weights were determined based on % Nitrogen (Table S1). MAYF source samples were consistently low in %N, therefore target weights were adjusted accordingly (Table S1). We aimed to obtain 3 replicates from each sample, though there was not always enough material to do so.

### Sample preparation of invertebrates for stable isotope analysis

Macroinvertebrates were identified to genus or lowest possible taxonomic level and assigned a FFG according to the Freshwater Insects CONUS trait database (Twardochleb *et al*. 2021). For individuals at the family-level, FFG was assigned based on the majority classification of genera nested within the family that were available in the database. Taxa not listed in the CONUS database were assigned a FFG based on alternative literature (Table S2). Chironomids were separated into Tanypodinae and Non-Tanypodinae. All chironomids were assigned to collector-gatherer as this is this most common feeding group for Chironomidae, aside from Tanypodinae which are predators (Berg 1995). When taxa had multiple feeding groups, we used the primary feeding group assigned. Macroinvertebrates were lyophilized in cryovials for 24-48 hours. Mollusk, gastropod, and crustacean tissues were removed from shells to avoid bias from inorganic carbon (Hill & McQuaid 2011). Amphipods and isopods were excluded due to inability to separate chiton from muscle tissue. Finally, predators were removed from the entire dataset to examine primary consumer diets only.

For each taxon, we aimed for 3 replicate samples. For larger taxa (≥ 0.800 mg), one sample was taken from each individual. For smaller taxa (i.e. midges, mayflies, etc.), multiple individuals were pooled into one sample to reach the target weight. Taxa were only included if we were able to obtain at least one replicate. Samples were weighed and packed into 5 x 3.5 mm tin capsules (Elemental Microanalysis, Part #: D1002) with target weights based on %N (Table S1). All samples were packed and shipped for stable isotope analysis of δ^13^C and δ^15^N at the UC Davis Stable Isotope Facility. Additional details on stable isotope analysis methods are provided in the Supplementary Methods: *Stable isotope analysis*.

### Dietary proportions

All data analysis was conducted using R v4.5 and v4.6 (R Core Team 2025, 2026). We used MixSIAR Bayesian mixing models (Stock *et al*. 2020, 2018; Stock & Semmens 2016) to estimate dietary proportions of basal resources consumed by macroinvertebrates. Because FBOM contains a mixture of allochthonous and autochthonous carbon, and was isotopically ambiguous as a discrete source, it was excluded from data analysis (Bonin *et al*. 2000). We defined source pools a priori based on autochthonous versus allochthonous origin: bryophytes, filamentous algae, macrophytes, and epilithon were considered autochthonous and leaf litter was considered allochthonous (Figures S1-S3). Two sites (MAYF07 and SYCA06) did not have samples of autochthonous sources; therefore, the basin average of the source means, standard deviations, and number of samples were used for those sites in mixing models.

We used two structures of MixSIAR models. The first, a control for comparing models stratified with covariates stratified by basin, included basin as a fixed effect and site as a random effect nested within basin, and selecting residual and process errors (Stock *et al*. 2020, 2018; Stock & Semmens 2016). The second model structure included FFG as a fixed effect, and site as a random effect, also selecting residual and process errors (Stock *et al*. 2020, 2018; Stock & Semmens 2016). This FFG model was replicated using the following univariate continuous fixed effects: % canopy cover, wetted width (m), drainage area (km^2^), proportion of dry days, mean annual air temperature, total annual precipitation (mm), aridity index, and allochthonous source quality (average C:N ratio). MixSIAR does not allow for multiple continuous effects in its modeling framework (Stock *et al*. 2020).

We used uninformed Dirichlet priors for all models (Stock *et al*. 2018). Trophic enrichment factors (TEFs) used were Δ13C = 0.3 ± 0.14‰ and Δ15N = 2.1 ± 0.21‰ based on recommendations for consumers analyzed whole (McCutchan Jr *et al*. 2003). All models were run with a chain length of 3,000,000, a burn-in of 1,500,000, a thinning of 500, and 3 chains (i.e. “extreme” in MixSIAR). Process error and residual error were set to “TRUE”. Convergence was assessed using Gelman-Rubin diagnostics ( R^; Gelman & Rubin 1992; Stock *et al*. 2018). All models exhibited acceptable convergence (maximum R^ = 1.09; no parameters exceeded R^ = 1.1). A small number of individual-level parameters exceeded the more stringent threshold of R^ = 1.05 in Models 7 and 10, but convergence was considered acceptable for model comparison and ecological inference (Stock & Semmens 2016; Table 2). We used the “loo” R package (Vehtari *et al*. 2017, 2026) to compare model fit leave-one-out cross validation (LOO) with a modified MixSIAR compare_models() function (Table 2; Stock *et al*. 2020). A small number of observations exhibited a Pareto-*k* value of (*k* > 1), therefore a sensitivity analysis was conducted to determine whether relative model support was influenced by observations with high Pareto-*k* values (See Supplementary Methods: *Model Comparison*; Tables S3 & S4).

**Table 2.** Model comparison using leave-one-out cross-validation information criterion (LOOic). Models are ranked by predictive performance, with lower LOOic indicating a better fit. *P*_diet_ is the estimated dietary proportion of consumers. pD is the effective number of parameters, se_lOOic is the standard error of LOOic and dLOOic is the difference from the best-performing model as well as the standard errors (se_dLOOic). Model weights show the relative support for each model.

| Fitness Rank | Model | Formula | pD | LOOic | se_LOOic | dLOOic | se_dLOOic | weight |
| --- | --- | --- | --- | --- | --- | --- | --- | --- |
| 1 | M7* | $P_{\text{diet}} \sim \text{FFG} + \text{Temperature} + (1 \mid \text{Site})$ | 450.9 | 2525.3 | 71.6 | 0.0 | NA | 0.389 |
| 2 | M9 | $P_{\text{diet}} \sim \text{FFG} + \text{Aridity Index} + (1 \mid \text{Site})$ | 447.8 | 2526.7 | 71.5 | 1.4 | 2.1 | 0.193 |
| 3 | M10* | $P_{\text{diet}} \sim \text{FFG} + \text{CN Ratio} + (1 \mid \text{Site})$ | 464.0 | 2527.4 | 72.1 | 2.1 | 1.6 | 0.136 |
| 4 | M8 | $P_{\text{diet}} \sim \text{FFG} + \text{Precipitation} + (1 \mid \text{Site})$ | 462.9 | 2528.1 | 71.5 | 2.8 | 2.4 | 0.096 |
| 5 | M5 | $P_{\text{diet}} \sim \text{FFG} + \text{Proportion of Dry Days} + (1 \mid \text{Site})$ | 459.7 | 2529.3 | 71.3 | 4.0 | 2.8 | 0.053 |
| 6 | M4 | $P_{\text{diet}} \sim \text{FFG} + \text{Wetted Width} + (1 \mid \text{Site})$ | 461.5 | 2529.7 | 71.9 | 4.4 | 2.1 | 0.043 |
| 7 | M2 | $P_{\text{diet}} \sim \text{FFG} + (1 \mid \text{Site})$ | 448.1 | 2530.5 | 72.0 | 5.2 | 2.5 | 0.029 |
| 8 | M3 | $P_{\text{diet}} \sim \text{FFG} + \text{Canopy} + (1 \mid \text{Site})$ | 469.1 | 2530.8 | 71.2 | 5.5 | 3.4 | 0.025 |
| 9 | M1 | $P_{\text{diet}} \sim \text{Basin} + (1 \mid \text{Basin/Site})$ | 447.9 | 2530.9 | 72.1 | 5.6 | 5.6 | 0.024 |
| 10 | M6 | $P_{\text{diet}} \sim \text{FFG} + \text{Drainage Area} + (1 \mid \text{Site})$ | 441.0 | 2532.0 | 72.6 | 6.7 | 3.4 | 0.014 |

\*Convergence assessed using Gelman–Rubin diagnostics ( $\hat{R}$ ). Models with parameters exceeding 1.05 are flagged; no model contained parameters with $\hat{R} > 1.10$ .

## RESULTS

### Environmental variables

Our basins, ranging two orders of magnitude in drainage area, also showed a wide range of climatic and hydrological conditions (Table 1). We found several significant correlations between environmental variables, with the two strongest being aridity index and precipitation (*r* = 0.976, p = 0.00) and C:N ratio and temperature (*r* = -0.65, p = 2.58 × 10⁻⁷, Figure 2). Further description can be found in Supplementary Results: *Summary of environmental variables*.

### Stable isotope values of sources and consumers

Average δ^13^C of source groups were similar, but average δ^15^N differed (Figure S4). Across basins, BLUE, KING, MAYF, and WPLP showed allochthonous sources being more depleted in δ^13^C, while PINN and SYCA allochthonous sources were more enriched in δ^13^C (Figure S4). Across all sites, δ^15^N was consistently lower for allochthonous sources than autochthonous (Figure S4).

All FFGs had similar average δ^13^C values (Figure S5). Average δ^15^N was similar across three of the four feeding groups, with shredders being relatively less enriched than the rest (Figure S5). Across basins, average δ^13^C of consumers was consistent, with WPLP being slightly more enriched and SYCA being slightly more depleted (Figure S5). Average δ^15^N of consumers displayed a broader range across sites with BLUE being the most enriched and WPLP being the least enriched (Figure S5).

### Relative proportions of allochthonous vs. autochthonous sources

Bayesian mixing model relative weights indicated that the top five best fit models were temperature (0.389), aridity index (0.193), C:N ratio of leaf litter (0.136), precipitation (0.096), and proportion of dry days (0.053, Table 2). For stream size variables measured, allochthony decreased with wetted width and drainage area and increased with canopy cover (Figures 3, S11, S15, S13). Climate variables showed a decrease in allochthony with temperature and a slight increase in allochthony with aridity and precipitation, with significant overlap of credible intervals (Figures 3, S6, S7, S9). Additionally, allochthony was positively associated with proportion of dry days (Figure 3, S10). C:N ratio showed a strong positive association with allochthony (Figures 3, S8). Dietary proportions were largely similar across FFGs, with shredders relying slightly more on allochthonous carbon (Figures 3, S12). At the basin level, SYCA and WPLP consumers consistently relied more on autochthonous carbon than other basins, with MAYF and PINN often also showing a slight dominance in autochthonous contributions (Figure S14). BLUE and KING were consistently dominated by allochthonous contributions (Figure S14). The basin-level model was outperformed by all other models, aside from drainage area (Table 2).

**Figure 3.**
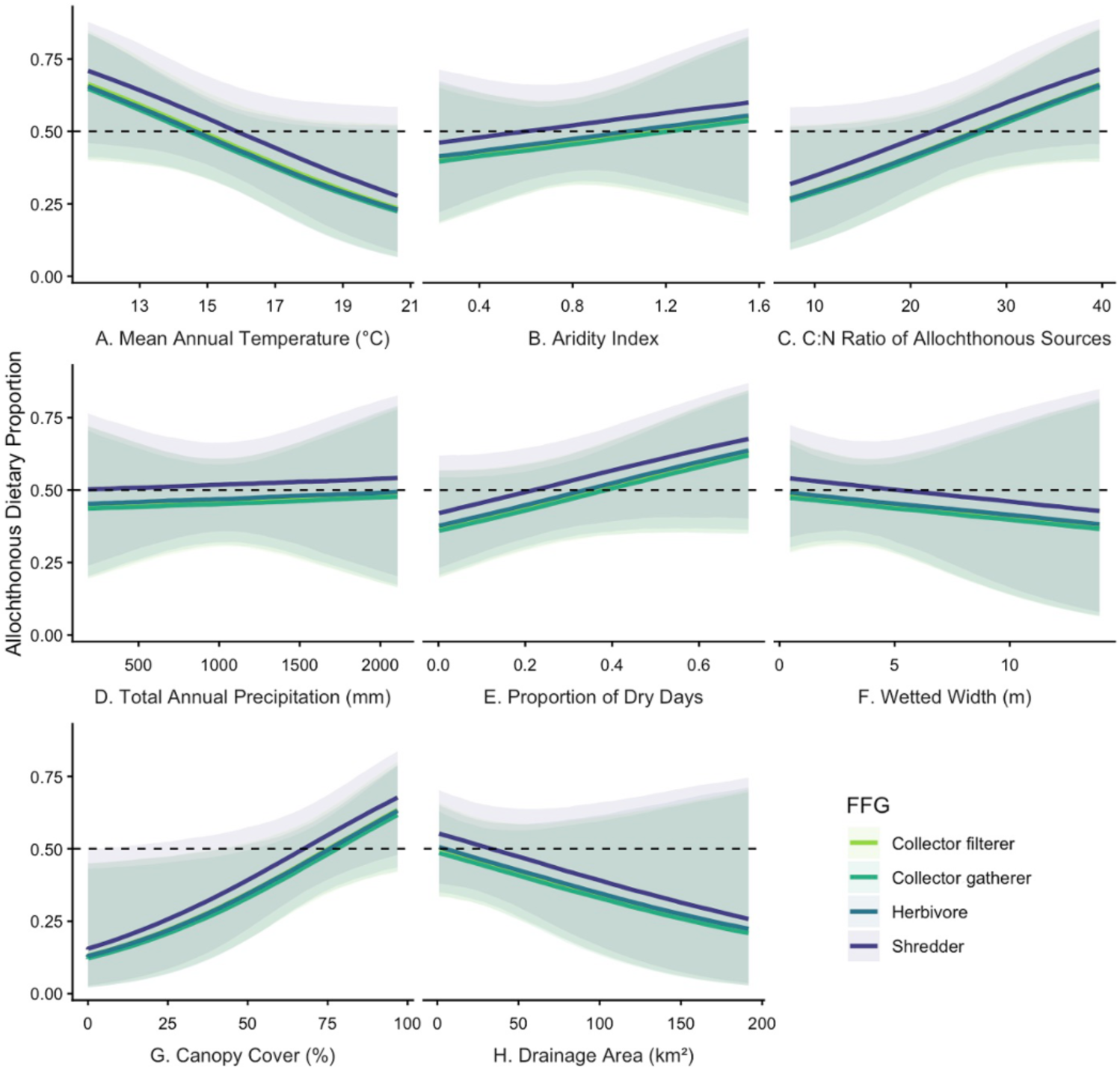
Posterior distribution with 95% credible intervals of consumer allochthonous dietary proportions, plotted against environmental covariates and colored by functional feeding group. Panels A-H are ordered by decreasing model weight, from left to right. Dashed lines represent 50% of the diet.

## DISCUSSION

Understanding mechanisms driving allochthony patterns in streams is critical to predicting ecosystem responses to changes in climate. Our findings suggest that invertebrates in smaller streams, and in cooler climates, likely depend more strongly on allochthonous inputs. All FFGs showed decreased allochthonous consumption with increasing temperature, with shredders shifting to an autochthonous-dominant diet at slightly higher temperatures than the rest. This pattern held true for wetted width and drainage area, yet uncertainty increased with stream size. This is likely because stream size was not evenly distributed across basins. Additionally, allochthony increased for all feeding groups with increasing C:N ratio, proportion of dry days, and canopy cover. Allochthony was largely unchanged with precipitation and aridity index. These results are consistent with our predictions and reinforce that the primary basal energy source in cool, non-perennial and headwater streams is allochthonous carbon.

### Influence of climate

Air temperature is an important driver of invertebrate resource consumption patterns at the local scale, as changes in air temperature are generally reflected by water temperature (Junker & Cross 2014; Morrill *et al*. 2005), but our study is the first to directly show an association between air temperature variation and allochthony in stream macroinvertebrates across a broad spatial scale. Our results indicate that collector-filterers, collector-gatherers, and herbivores shift to a predominantly autochthonous diet when mean annual air temperature reaches approximately 15 °C, with a slightly higher threshold for shredders. Air temperature is indirectly linked to stream primary production, as air temperature drives stream temperature (Patrick *et al*. 2019). Increasing stream temperatures are linked to higher metabolic demands and consumption for macroinvertebrates (Bonacina *et al*. 2023), meaning macroinvertebrates in warmer streams may require higher quality food sources like algae. Warmer streams are shown to have higher primary production (Demars *et al*. 2011; Hood *et al*. 2018; Morin *et al*. 1999), increasing the availability of autochthonous food sources. Therefore, a possible explanation for the shift to an autochthonous-dominant diet is preference. While our aridity index and precipitation models showed less profound correlations with allochthony than expected, both models show shredders shifting to allochthonous diets as aridity index and precipitation increase. The aridity index model was the next best fit of the climate variables; however, we observed strong collinearity between precipitation and aridity index, therefore it is likely that neither is more ecologically relevant than the other. Because aridity index accounts for precipitation and evapotranspiration, aridity index may better capture long-term, climate-driven water availability. Extreme aridity can also facilitate stream drying, which has been shown to decrease algal biomass (Nelson *et al*. 2023). Therefore, the slight shift to allochthonous food sources with aridity may be a function of resource limitation. These results, particularly of the temperature model, reflect the idea that metabolic compensation likely occurs when resource availability changes in streams (Dodds *et al*. 2015, 2019). In other words, energetic contributions in streams may switch between terrestrial carbon and primary production depending on preference and availability.

### Influence of streamflow permanence

Studies have shown mixed and often contrasting effects of stream drying on allochthony. Some research has found that aquatic invertebrates assimilate more autochthonous carbon as streamflow intermittency increases (Dekar *et al*. 2009; Siebers *et al*. 2019). However, a global scale meta-analysis has shown greater allochthony for macroinvertebrates in arid climates (Allen *et al*. 2024). Here, we found that autochthonous inputs dominated in Sycamore Creek, AZ, an arid, desert stream system with many non-perennial reaches. This is likely due to the combination of a lack of canopy cover and increased light availability within this watershed. Additionally, Sycamore Creek has been shown to be predominantly fueled by autochthonous carbon, apart from post-flooding events, where algae tend to be scoured away from high volumes of water and terrestrial detritus is transported downstream (Fisher *et al*. 1982). We also found that the proportion of dry days model had a lower predictive performance than the aridity index model. Aridity influences terrestrial carbon uptake (Zhou *et al*. 2019); therefore, it may limit plant growth (i.e. allochthonous inputs). Our model showed that allochthony increases with the proportion of dry days, so it is likely that the dominance of autochthonous sources in Sycamore Creek is a function of other variables, like temperature and aridity, in combination with drying.

### Influence of stream size

Stream size directly influences allochthony, and our models indicate that macroinvertebrates consume more autochthonous carbon as stream width increases. This relationship supports the idea that as streams grow from first to mid-order, it relies more on primary production (Vannote *et al*. 1980). We also found an increase in allochthony relative to canopy cover, confirming that smaller streams with more canopy cover rely more on terrestrial detritus as a primary food source. Drainage area was associated with decreased allochthony with considerably large uncertainty and was the worst predictor. This is likely because wetted width is a direct measure of the stream morphology that has been shown to influence within stream gross primary production by controlling light input (Bernhardt *et al*. 2022). Canopy cover, a direct measurement of the surrounding riparian environment, is indicative of local allochthonous carbon inputs. Contrastingly, drainage area is not a direct measurement of local conditions, but a proxy for stream size that does not include channel morphology or other potentially biologically relevant features of the stream habitat.

### Influence of biological traits

Increased C:N ratio is indicative of detritus that is rich in carbon and poor in protein content or nitrogen, making it a lower quality food source (Lau *et al*. 2009). Macroinvertebrates have been shown to prefer leaves with lower C:N ratios (König *et al*. 2014), due to the higher nutritional value. Our findings contradict our hypothesis that allochthony will increase with lower C:N ratios (i.e. higher nutritional value). Instead, we found that allochthony increased with decreasing leaf litter quality. This finding is accompanied by a moderate collinearity between temperature and C:N ratio, showing that warmer streams had leaf litter with lower C:N ratios, and thus were a higher quality food resource. This trend has been observed globally across varying climates through rapid breakdown of C content (de la Casa *et al*. 2025). However, warmer streams also have higher primary production (Demars *et al*. 2011; Hood *et al*. 2018; Morin *et al*. 1999); therefore, it is possible that even when autochthonous sources were lower in C:N, autochthonous sources were still more available and preferred. Ultimately, we interpret our results to indicate that C:N ratio is not a determining factor of dietary patterns in our study, but rather a secondary effect of temperature.

Functional feeding traits have been shown to explain variation in allochthonous contributions (Allen *et al*. 2024). Our results show that shredders consume more allochthonous carbon than other FFGs, aligning with our hypothesis that consumer diets would be shaped by FFG. However, differences in dietary proportions between collector-gatherers, collector-filterers, and herbivores were negligible. This suggests that resource limitation might be more important in streams, such that feeding traits becomes less meaningful when there are so few options due to disturbance or environmental extremes. Macroinvertebrates can feed selectively in response to environmental change (Guo *et al*. 2018), therefore, feeding groups may shift their food sources according to availability determined by the effects of climate, drying, and stream size. Furthermore, it is possible that some consumers are passively consuming detrital carbon when seeking out the microbial and algal communities present on leaves as detrital material can comprise 30% of periphyton (Rasmussen 2010). In other words, in a perennial stream, a scraper may be feeding on epilithon on a rock but in periods of drying, they may result to scraping the periphyton off leaves, in turn consuming untargeted allochthonous material, if not consuming the leaf material itself. Our results show that FFG alone is not the most important driver of allochthony and suggest that FFG classifications may not always be an accurate reflection of macroinvertebrate diets, particularly in resource-limited systems.

### Limitations and considerations

There are several limitations that should be considered when contextualizing these findings. Firstly, MixSIAR does not allow for much flexibility when developing models, such as the ability to only model two categorical effects and one continuous effect at a time. Therefore, we acknowledge that there may be interactions between the several environmental variables tested that were not able to test in our modeling framework, in addition to other environmental variables not addressed in this study. This highlights the need for a more comprehensive modeling approach for stable isotope mixing models that are more compatible with complex datasets and can incorporate several continuous effects, while maintaining a hierarchical structure.

Another limitation is that there are several ways to quantify stream drying, and in this study, we have selected one metric to be representative of flow characterization. While other metrics exist besides annual proportion of dry days, this metric has been used comprehensively to compare several biodiversity studies (Datry *et al*. 2014). In addition to the environmental variables compared in this study, there are likely other important factors to consider that may be difficult to extrapolate to a large spatial scale. Additionally, stable isotope analyses also carry several inherent challenges such as poor source discrimination and limitation by the number of tracers used (Stock *et al*. 2018). The addition of a third tracer, such as δ^2^H or δ^34^S could greatly improve source discrimination (Doucett *et al*. 2007; Raoult *et al*. 2024), reducing overlap in isotopic space of sources. Lastly, the assignment of a singular FFG is limiting, as many taxa show characteristics of several feeding groups, however, we believe that we have provided the best representation of each taxon’s functional role in streams.

### Conclusion

This is the first study, to our knowledge, to empirically investigate which factors are associated with basal resource contributions in stream food webs across a sub-continental scale. Our results show that macroinvertebrate dietary proportions are linked primarily to changes temperature, but that flow permanence, stream size, and consumer biological traits are also important. We propose that the observed response of macroinvertebrate feeding habits is a function of resource limitation, however, it is possible that environmental stressors on invertebrates such as thermal or drought stress changes feeding behavior (Aspin *et al*. 2019; Vannote & Sweeney 1980). We also suggest that future work incorporates system hydrology and climate effects when investigating food webs in streams. As global temperatures increase, streams are predicted to become warmer, thus increasing oxygen stress of stream organisms (Sand-Jensen & Pedersen 2005). Here we show that climate, specifically air temperature, is important for predicting energy sources in streams, deeming this work increasingly important for developing a unifying theory to predict nutrient cycling and energy flow in these systems under global change.

## ACKNOWLEDGEMENTS

This work was supported by the US National Science Foundation awards 2207680, 2150626, 1802766, 1802714, 1802811; and the USDA National Institute of Food and Agriculture and Hatch Appropriations under Project #PEN04817 and Accession #7003724.

