## Supporting Information for "Allochthony in stream food webs decreases with temperature across a subcontinental scale"

Summary: The supporting information includes supplementary methods, supplementary results, 3 tables, 15 figures, and references.

**Supplementary Methods**

*Stable isotope analysis*

Stable isotope analysis followed the UC Davis Stable Isotope Facility protocol (Brandeberry 2020). Invertebrate and basal resource samples were combusted using a PDZ Europa ANCA-GSL elemental analyzer at 950°C in a reactor packed with chromium oxide and silvered copper oxide to ensure complete oxidation. The oxides were then eliminated using a reduction reactor (reduced copper at 650°C). Helium was used to transport gases through a water trap containing magnesium perchlorate and phosphorus pentoxide, which remove water vapor from the gases. Finally, gases were separated into nitrogen and carbon dioxide based on retention times, using a gas chromatography column (65°C, 65 mL/min), and transferred to a PDZ Europa 20-20 isotope ratio mass spectrometer (Sercon Ltd., Cheshire, UK). For the portion of samples that were collected on GFFs, either an Elementar Vario El Cube or Micro Cube elemental analyzer (Elementar Analysensysteme GmbH, Hanau, Germany) was used to combust at 1080°C using the same protocol. The carbon dioxide was kept in an absorption trap until the nitrogen was analyzed. It was then heated and the carbon dioxide was released to either an Isoprime VisION IRMS (Elementar Uk Ltd, Cheadle, UK) or a PDZ Europa 20-20 isotope ratio mass spectrometer (Sercon Ltd., Cheshire, UK).

The samples were calibrated against the following international standards for D13C and D15N: IAEA-600, USGS40, USGS41, USGS42, USGS43, USGS61, USGS64, and USGS65. Additionally, the following in-house QA/QC reference materials were used for instrument calibration: alfalfa flour, amaranth flour, caffeine, chitin, enriched alanine, glutamic acid, glutathione, scallop, and nylon powder. Sample initial isotope ratios were adjusted based on known reference material values.

*Model Comparison*

We compared candidate models using leave-one-out cross validation (LOO). Because the compare_models() function in MixSIAR ( v3.1.12; Stock *et al.* 2020, 2018; Stock & Semmens 2016) was not fully compatible with the output structure of loo ( v.2.10.1; Vehtari *et al.* 2017, 2026), we modified the function, while retaining the original MixSIAR approach to calculating LOOIC and model weights. We used Pareto-*k* estimates for each model to evaluate lo diagnostics. A small number of observations (obs. 66, 143, and 390) had a Pareto-*k* value of $k > 1$ (Table S3). To evaluate the sensitivity of the model comparison to the high Pareto-*k* values, we conducted a sensitivity analysis by conducting the model comparison without the problematic observations. Models were not refit; therefore, the sensitivity analysis addresses the sensitivity of predictive model comparison to these observations rather than the effect of removing the observations. We found ΔLOOIC remained within 4 units across models and model weights (Table 2, Table S4) and the model with the most predictive support remained consistent. While the 2$^{nd}$ and 3$^{rd}$ ranked models swapped, they both remained in the top 3 with the best predictive support. Therefore, we elected to use the full LOO data for model comparison.

**Supplementary Results**

*Summary of environmental variables*

Across all basins, SYCA had the largest drainage area (57.07 ± 70.85$km^2$), while WPLP had the smallest (0.57 ± 0.62$km^2$, Table 1). PINN had the smallest wetted width (1.93 ± 1.04m) and MAYF had the largest (4.09 ± 3.67m, Table 1). Canopy cover was lowest at PINN (37.4 ± 33.34%) and highest at WPLP (85.86 ± 9.48%, Table 1).

WPLP also experienced the most precipitation (1808.72 ± 222.22 mm) and highest aridity index, indicating the least aridity (1.28 ± 0.12, Table 1). PINN was the most arid basin, with the lowest annual precipitation (217.78 ± 12.11 mm) and the lowest aridity index (0.27 ± 0.05, Table 1). KING had the lowest mean annual temperature (12.39 ± 0.08°C), while MAYF had the highest (18.24 ± 0.06 °C, Table 1).

In addition to temperature, MAYF was also the most perennial, having the lowest proportion of dry days (0.01 ± 0.02, Table 1). BLUE was the most intermittent, with the highest proportion of dry days (0.41 ± 0.20, Table 1). Allochthonous sources at WPLP had the highest C:N ratio (30.98 ± 4.44, Table 1), indicative of the lowest nutritional value. MAYF allochthonous sources had the lowest C:N ratio (13.62 ± 1.23, Table 1), indicative of the highest nutritional value. Results of a Pearson pairwise correlation test showed several significant correlations between environmental variables, with the two strongest being aridity index and precipitation ($r = 0.976$, $p = 0.00$) and C:N ratio and temperature ($r = -0.65$, $p = 2.58 \times 10^{-7}$, Figure 2).

**Table S1.** Target weights for each sample type. MAYF sources are distinguished from the rest of the basins as target weights were increased to adjust for low %N in those samples.

| **Sample Type** | **Tin Size** | **Target Weight (mg)** | **Range (∓10%)** | **Est. %N** | **Target N weight (ug)** |
|---|---|---|---|---|---|
| Leaf litter | 9 x 5 mm | 8.000 | 7.2-8.8 | 1% | 80 |
| Bryophytes | 9 x 5 mm | 8.000 | 7.2-8.8 | 1% | 80 |
| FBOM | 9 x 5 mm | 20.000 | 18-22 | 0.4% | 80 |
| Epilithon | 9 x 5 mm | 10.000 | 9-11 | 0.75% | 80 |
| Macrophytes | 9 x 5 mm | 4.000 | 3.6-4.4 | 2% | 80 |
| Filamentous algae | 9 x 5 mm | 5.250 | 4.725-5.775 | 1.5% | 80 |
| Invertebrates | 5 x 3.5 mm | 0.800 | 0.72-0.88 | 10% | 80 |
| Leaf litter (MAYF) | 9 x 5 mm | 8.000 | 7.2-8.8 | 1% | 80 |
| FBOM (MAYF) | 9 x 5 mm | 20.000 | 18-22 | 0.05% | 80 |
| Epilithon (MAYF) | 9 x 5 mm | 20.000 | 18-22 | 0.175% | 80 |
| Macrophytes (MAYF) | 9 x 5 mm | 8.000 | 7.2-8.8 | 0.175% | 80 |
| Filamentous algae (MAYF) | 9 x 5 mm | 20.000 | 18-22 | ~1% | 80 |

**Table S2.** Table of all taxa identified, prior to removal of predators, labeled as the lowest taxonomic ID in alphabetical order, and the associated functional feeding group and the source used to validate the assignment.

| **Lowest ID** | **Feeding Group** | **Source** |
|---|---|---|
| *Abedus* | Predator | (Twardochleb *et al.* 2021) |
| *Acroneuria* | Predator | (Twardochleb *et al.* 2021) |
| *Agabinus* | Predator | (Twardochleb *et al.* 2021) |
| *Agabus* | Predator | (Twardochleb *et al.* 2021) |
| *Agabus_adult* | Predator | (Twardochleb *et al.* 2021) |
| *Agabus_larvae* | Predator | (Twardochleb *et al.* 2021) |
| *Agnetina* | Predator | (Twardochleb *et al.* 2021) |
| *Alloperla* | Collector.gatherer | (Twardochleb *et al.* 2021) |
| *Ameletus* | Collector.gatherer | (Twardochleb *et al.* 2021) |
| *Amphinemura* | Shredder | (Twardochleb *et al.* 2021) |
| *Anchytarsus* | Shredder | (Twardochleb *et al.* 2021) |
| *Angarotipula* | Collector.gatherer | (Yang *et al.* 2022) |
| Anisoptera | Predator | (Sentis *et al.* 2022) |
| *Antocha* | Collector.gatherer | (Twardochleb *et al.* 2021) |
| *Aquarius* | Predator | (Twardochleb *et al.* 2021) |
| *Aquarius_adult* | Predator | (Twardochleb *et al.* 2021) |
| *Argia* | Predator | (Twardochleb *et al.* 2021) |
| *Arigomphus* | Predator | (Twardochleb *et al.* 2021) |
| Baetidae | Collector.gatherer | (Twardochleb *et al.* 2021), used genus level |
| *Belostoma* | Predator | (Twardochleb *et al.* 2021) |
| *Berosus* | Herbivore | (Twardochleb *et al.* 2021) |
| *Berosus_adult* | Herbivore | (Twardochleb *et al.* 2021) |
| *Berosus_larvae* | Herbivore | (Twardochleb *et al.* 2021) |
| *Berosus_salvini* | Herbivore | (Twardochleb *et al.* 2021) |
| *Boreonectes* | Predator | (Yee 2023) |
| *Boyeria* | Predator | (Twardochleb *et al.* 2021) |
| *Buenoa* | Predator | (Twardochleb *et al.* 2021) |
| *Caenis* | Collector.gatherer | (Twardochleb *et al.* 2021) |
| *Caloparyphus* | Collector.gatherer | (Twardochleb *et al.* 2021) |
| *Calopteryx* | Predator | (Twardochleb *et al.* 2021) |
| *Cambarus* | Shredder | (Helms & Creed 2005) |

| | | |
|---|---|---|
| *Capnia* | Shredder | (Twardochleb *et al.* 2021) |
| Capniidae | Shredder | (Twardochleb *et al.* 2021), used genus level |
| *Celina_adult* | Predator | (Twardochleb *et al.* 2021) |
| *Celina_larvae* | Predator | (Twardochleb *et al.* 2021) |
| *Cheumatopsyche* | Collector.filterer | (Twardochleb *et al.* 2021) |
| Chironomidae | Collector.gatherer | (Twardochleb *et al.* 2021), used genus level, (Berg 1995) |
| *Chironomus* | Collector.gatherer | (Twardochleb *et al.* 2021) |
| Chloroperlidae | Predator | (Majdi *et al.* 2015), (Twardochleb *et al.* 2021), used genus level |
| *Copelatus* | Predator | (Twardochleb *et al.* 2021) |
| *Corbicula* | Collector.filterer | (Cohen *et al.* 1984) |
| *Cordulegaster* | Predator | (Twardochleb *et al.* 2021) |
| *Corixini* | Collector.gatherer | (Sutton 1951) |
| Crambidae | Shredder | (De-Freitas *et al.* 2019) |
| *Cryphocricos* | Predator | (Twardochleb *et al.* 2021) |
| *Culex* | Collector.filterer | (Twardochleb *et al.* 2021) |
| Culicidae_pupae | Collector.filterer | (Twardochleb *et al.* 2021), used genus level |
| *Dineutus* | Predator | (Twardochleb *et al.* 2021) |
| *Diplectrona* | Collector.filterer | (Twardochleb *et al.* 2021) |
| *Dubiraphia* | Collector.gatherer | (Twardochleb *et al.* 2021) |
| Dysticidae | Predator | (Miller & Bergsten 2023), (Twardochleb *et al.* 2021), used genus level |
| Dytiscinae | Predator | (Miller and Bergsten 2023), (Twardochleb *et al.* 2021), used genus level |
| *Eccoptura* | Predator | (Twardochleb *et al.* 2021) |
| *Epeorus* | Collector.gatherer | (Twardochleb *et al.* 2021) |
| *Ephemerella* | Collector.gatherer | (Twardochleb *et al.* 2021) |
| *Euparyphus* | Collector.gatherer | (Twardochleb *et al.* 2021) |
| *Eurylophella* | Collector.gatherer | (Twardochleb *et al.* 2021) |
| *Fallceon* | Collector.gatherer | (Twardochleb *et al.* 2021) |
| *Faxonius* | Collector.gatherer | (Helms & Creed 2005) |
| *Gerris* | Predator | (Twardochleb *et al.* 2021) |
| *Gyraulus* | Herbivore | (Dillon 2006) |
| *Gyretes* | Predator | (Twardochleb *et al.* 2021) |
| *Gyrinus* | Predator | (Twardochleb *et al.* 2021) |
| *Hagenius_brevistylus* | Predator | (Twardochleb *et al.* 2021) |
| *Haliplus* | Herbivore | (Twardochleb *et al.* 2021) |
| *Haploperla* | Predator | (Twardochleb *et al.* 2021) |
| *Helichus* | Herbivore | (Twardochleb *et al.* 2021) |

| | | |
|---|---|---|
| *Helichus_adult* | Herbivore | (Twardochleb *et al.* 2021) |
| *Hemerodromia* | Predator | (Twardochleb *et al.* 2021) |
| *Hetaerina* | Predator | (Twardochleb *et al.* 2021) |
| *Hexatoma* | Predator | (Twardochleb *et al.* 2021) |
| *Homoplectra* | Collector.filterer | (Twardochleb *et al.* 2021) |
| *Hydatophylax* | Shredder | (Twardochleb *et al.* 2021) |
| Hydrachnidiae | Predator | (Di Sabatino *et al.* 2000) |
| Hydroporinae | Predator | (Longing *et al.* 2017), (Twardochleb *et al.* 2021), used genus level |
| Hydroporinae_larvae | Predator | (Longing *et al.* 2017), (Twardochleb *et al.* 2021), used genus level |
| *Hydropsyche* | Collector.filterer | (Twardochleb *et al.* 2021) |
| *Hygrotus* | Predator | (Twardochleb *et al.* 2021) |
| *Illybius* | Predator | (O'Connor *et al.* 2021) |
| *Ischnura* | Predator | (Twardochleb *et al.* 2021) |
| *Isonychia* | Collector.filterer | (Twardochleb *et al.* 2021) |
| *Isoperla* | Predator | (Twardochleb *et al.* 2021) |
| *Laccobius* | Herbivore | (Twardochleb *et al.* 2021) |
| *Laccobius_adult* | Herbivore | (Twardochleb *et al.* 2021) |
| *Laccobius_larvae* | Herbivore | (Twardochleb *et al.* 2021) |
| *Laccophilus_pictus* | Predator | (Twardochleb *et al.* 2021) |
| *Lanthus* | Predator | (Twardochleb *et al.* 2021) |
| *Lepidostoma* | Shredder | (Twardochleb *et al.* 2021) |
| Leptophlebiidae | Collector.gatherer | (Petersen & Cummins 1974) |
| *Lestes* | Predator | (Twardochleb *et al.* 2021) |
| *Libellula* | Predator | (Twardochleb *et al.* 2021) |
| *Limnephilus* | Shredder | (Twardochleb *et al.* 2021) |
| *Liodessus* | Predator | (Twardochleb *et al.* 2021) |
| *Liodessus_adult* | Predator | (Twardochleb *et al.* 2021) |
| *Lymnaea* | Hebrivore | (Baker 1911) |
| *Macronychus* | Collector.gatherer | (Twardochleb *et al.* 2021) |
| *Mesocapnia* | Shredder | (Twardochleb *et al.* 2021) |
| *Metrobates* | Predator | (Twardochleb *et al.* 2021) |
| *Microvelia* | Predator | (Twardochleb *et al.* 2021) |
| Naididae | Collector.gatherer | (Mariom & Mollah 2013) |
| *Nasiaeschna_pentacantha* | Predator | (Twardochleb *et al.* 2021) |
| *Neocorixa* | Collector.gatherer | (Sutton 1951) |
| *Neohermes* | Predator | (Twardochleb *et al.* 2021) |
| *Neophylax* | Herbivore | (Twardochleb *et al.* 2021) |
| *Neoporus* | Predator | (Twardochleb *et al.* 2021) |

| | | |
|---|---|---|
| *Neoporus_adult* | Predator | (Twardochleb *et al.* 2021) |
| *Neoporus_larvae* | Predator | (Twardochleb *et al.* 2021) |
| *Neurocordulia* | Predator | (Twardochleb *et al.* 2021) |
| *Nigronia* | Predator | (Twardochleb *et al.* 2021) |
| *Notonecta* | Predator | (Twardochleb *et al.* 2021) |
| *Notonecta_kirbyi* | Predator | (Twardochleb *et al.* 2021) |
| *Octogomphus* | Predator | (Twardochleb *et al.* 2021) |
| Oligochaeta | Collector.gatherer | (Mariom & Mollah 2013) |
| *Ophiogomphus* | Predator | (Twardochleb *et al.* 2021) |
| *Optioservus* | Herbivore | (Twardochleb *et al.* 2021) |
| *Optioservus_adult* | Herbivore | (Twardochleb *et al.* 2021) |
| *Optioservus_larvae* | Herbivore | (Twardochleb *et al.* 2021) |
| *Oxycera* | Herbivore | (Mángano *et al.* 1996) |
| *Paragnetina* | Predator | (Twardochleb *et al.* 2021) |
| *Paraleptophlebia* | Collector.gatherer | (Twardochleb *et al.* 2021) |
| *Pelonomus_obscurus* | Herbivore | (Twardochleb *et al.* 2021) |
| *Peltodytes* | Herbivore | (Twardochleb *et al.* 2021) |
| *Peltodytes_adult* | Herbivore | (Twardochleb *et al.* 2021) |
| *Peltodytes_larvae* | Herbivore | (Twardochleb *et al.* 2021) |
| *Perlesta* | Predator | (Twardochleb *et al.* 2021) |
| *Physa* | Herbivore | (Brown *et al.* 1994) |
| *Pisidium* | Collector.filterer | (Martin 1998) |
| *Planorbella* | Herbivore | (Van Horn *et al.* 2012) |
| *Pleurocera* | Herbivore | (Brown *et al.* 2008) |
| *Postelichus* | Herbivore | (Twardochleb *et al.* 2021) |
| *Progomphus* | Predator | (Twardochleb *et al.* 2021) |
| *Prosimulium* | Collector.filterer | (Twardochleb *et al.* 2021) |
| *Psephenus* | Herbivore | (Twardochleb *et al.* 2021) |
| *Psilotreta* | Herbivore | (Twardochleb *et al.* 2021) |
| *Pteronarcys* | Shredder | (Twardochleb *et al.* 2021) |
| *Remenus* | Predator | (Twardochleb *et al.* 2021) |
| *Rhagovelia* | Predator | (Twardochleb *et al.* 2021) |
| *Rhyacophila* | Predator | (Twardochleb *et al.* 2021) |
| *Sanfilippodytes* | Predator | (Twardochleb *et al.* 2021) |
| Schoenobiinae | Shredder | (De-Freitas *et al.* 2019) |
| Sciomyzidae | Predator | (Berg & Knutson 1978) |
| *Sialis* | Predator | (Twardochleb *et al.* 2021) |
| *Simulium* | Collector.filterer | (Twardochleb *et al.* 2021) |
| *Somatochlora* | Predator | (Twardochleb *et al.* 2021) |

| | | |
|---|---|---|
| *Sperchopsis_tessellata* | Predator | (Twardochleb *et al.* 2021) |
| *Stenelmis_adult* | Herbivore | (Twardochleb *et al.* 2021) |
| *Stenelmis_larvae* | Herbivore | (Twardochleb *et al.* 2021) |
| *Stenonema* | Herbivore | (Twardochleb *et al.* 2021) |
| *Stenonema_femoratum* | Herbivore | (Twardochleb *et al.* 2021) |
| *Stratiomys* | Collector.gatherer | (Twardochleb *et al.* 2021) |
| *Sympetrum* | Predator | (Twardochleb *et al.* 2021) |
| *Tabanus* | Predator | (Twardochleb *et al.* 2021) |
| *Tallaperla* | Shredder | (Twardochleb *et al.* 2021) |
| Tanypodinae | Predator | (Baker & McLachlan 1979) |
| *Thermonectus_basillaris* | Predator | (Twardochleb *et al.* 2021) |
| *Tipula* | Shredder | (Twardochleb *et al.* 2021) |
| Tipulidae | Shredder | (Twardochleb *et al.* 2021), used genus level |
| *Trepobates* | Predator | (Twardochleb *et al.* 2021) |
| *Trichocorixa* | Predator | (Twardochleb *et al.* 2021) |
| *Tricorythodes* | Collector.gatherer | (Twardochleb *et al.* 2021) |
| *Tropisternus_adult* | Predator | (Twardochleb *et al.* 2021) |
| *Uvarus* | Predator | (Twardochleb *et al.* 2021) |
| *Wormaldia* | Collector.filterer | (Twardochleb *et al.* 2021) |
| *Zoraena* | Predator | (Sentis *et al.* 2023) |

**Table S3.** Pareto-*k* diagnostics for candidate models, including the maximum Pareto-*k*, effective number of parameters ($P_D$) number of observations with k > 0.7 and k > 1.

| **model** | $P_D$ | **max_k** | **n_k_gt_0.7** | **n_k_gt_1** |
|---|---|---|---|---|
| basin_only | 447.9 | 1.27314806 | 7 | 2 |
| ffg_only | 448.1 | 1.20782395 | 8 | 2 |
| ffg_canopy | 469.1 | 1.08527701 | 6 | 2 |
| ffg_CN | 464.0 | 1.12684422 | 8 | 2 |
| ffg_aridity | 447.8 | 1.12798202 | 8 | 1 |
| ffg_drainage | 441.0 | 1.38120123 | 10 | 2 |
| ffg_dry | 459.7 | 1.03235632 | 8 | 1 |
| ffg_ppt | 462.9 | 1.26999718 | 10 | 2 |
| ffg_temp | 450.9 | 1.09898981 | 7 | 1 |
| ffg_wet | 461.5 | 1.16039807 | 8 | 1 |

**Table S4.** Sensitivity analysis of candidate models using leave-one-out cross-validation (LOO) after excluding observations with Pareto-$k > 1$.

| Model | LOOic | se_LOOic | dLOOic | se_dLOOic | weight |
|---|---|---|---|---|---|
| ffg_temp | 2458.6 | 63.8 | 0 | NA | 0.308 |
| ffg_CN | 2459.3 | 64 | 0.7 | 1.4 | 0.212 |
| ffg_aridity | 2460.3 | 63.8 | 1.7 | 2.1 | 0.131 |
| ffg_ppt | 2461.3 | 63.8 | 2.8 | 2.1 | 0.077 |
| ffg_drainage | 2461.5 | 63.7 | 2.9 | 2.1 | 0.072 |
| ffg_wet | 2462 | 63.8 | 3.4 | 2.1 | 0.055 |
| ffg_only | 2462.1 | 63.7 | 3.6 | 2.2 | 0.051 |
| basin_only | 2462.9 | 64 | 4.3 | 5.5 | 0.035 |
| ffg_dry | 2463.2 | 63.7 | 4.6 | 2.8 | 0.031 |
| ffg_canopy | 2463.4 | 63.4 | 4.8 | 2.7 | 0.028 |

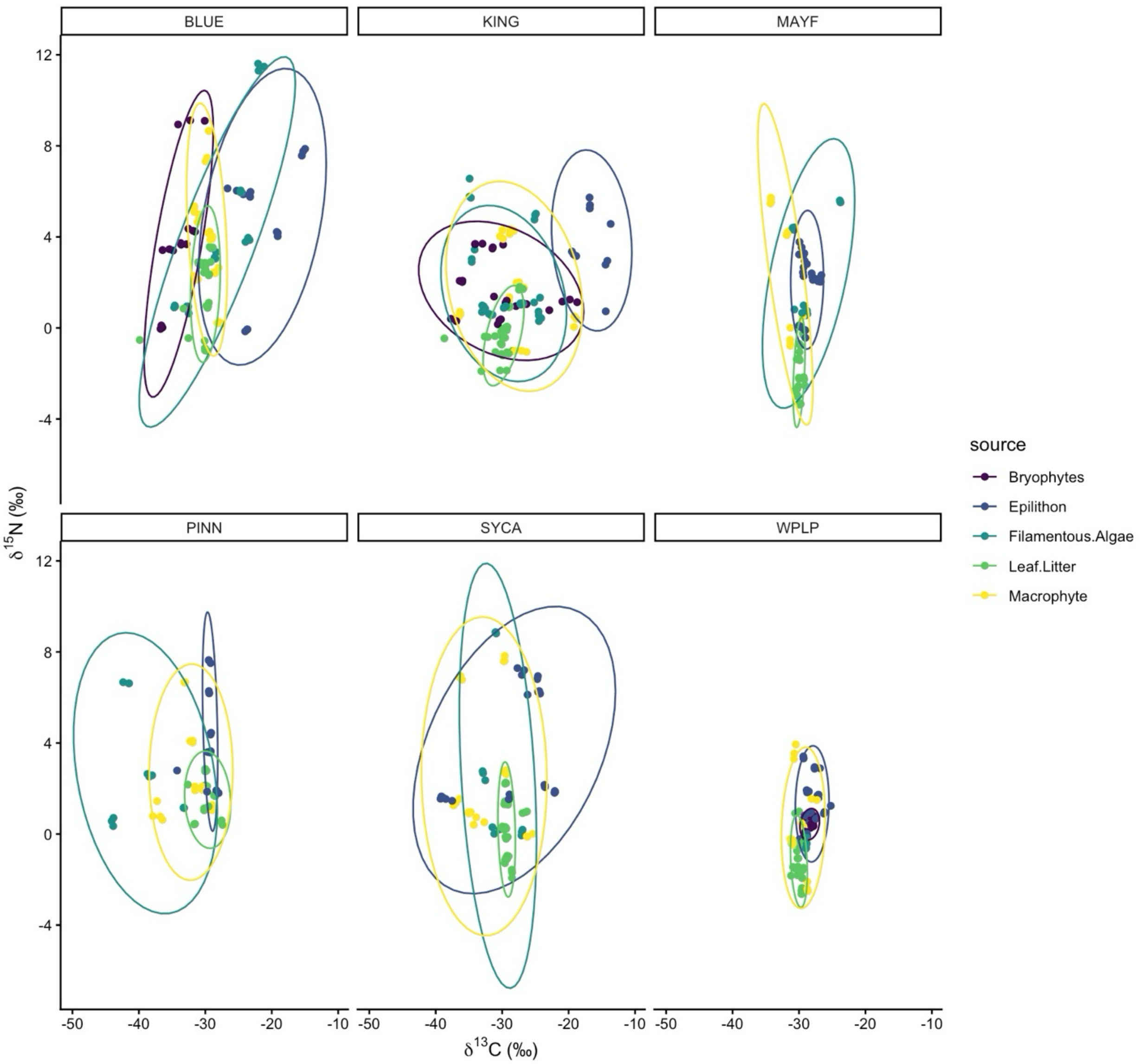


**Figure S1.** Isotopic bi-plot of ungrouped source data, colored by source type.

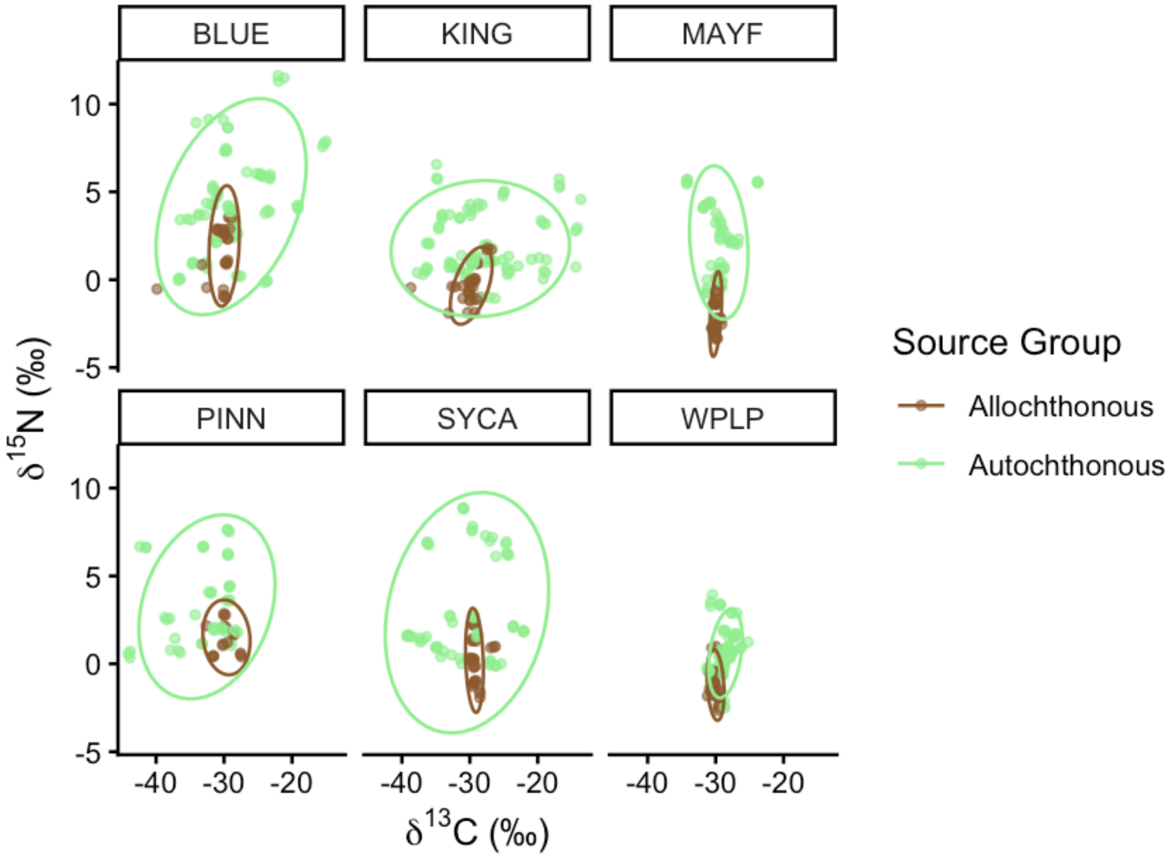


**Figure S2.** Isotopic bi-plot showing $\delta^{13}C$ and $\delta^{15}N$ values for source data, grouped and colored by source type.

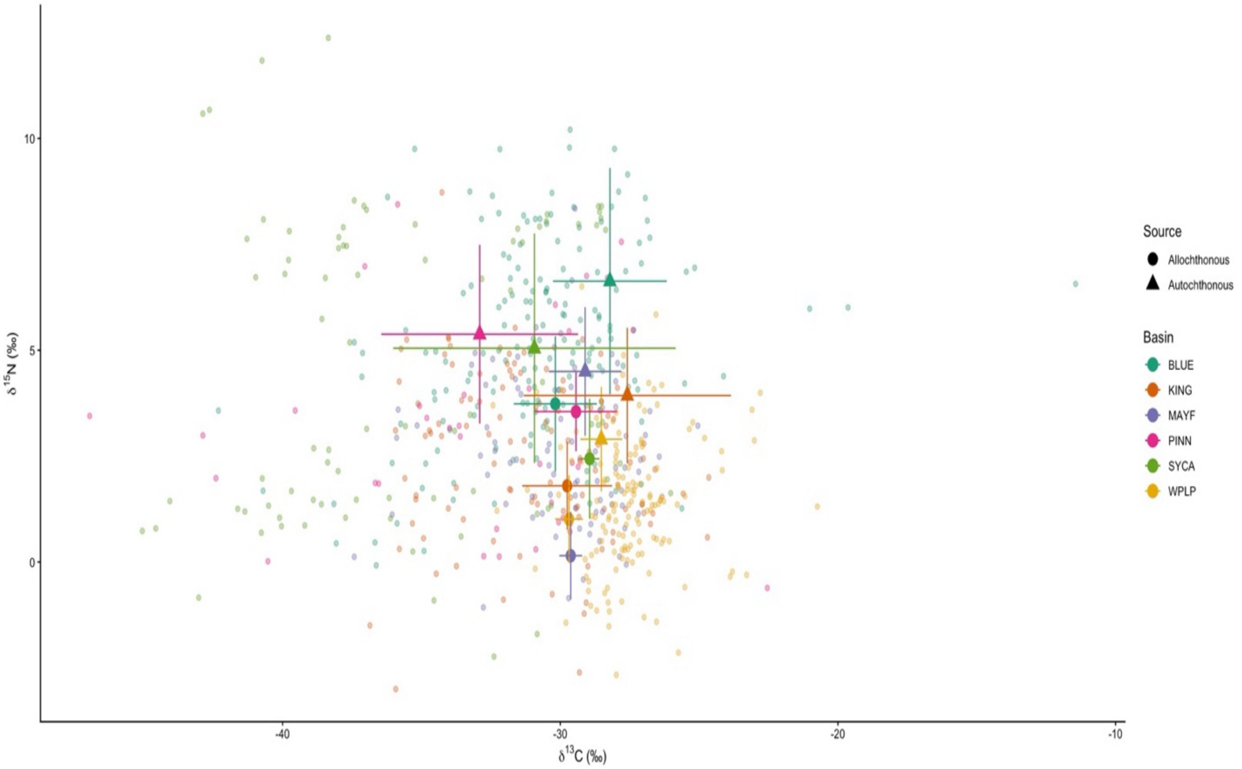


**Figure S3.** Isotopic bi-plot showing $\delta^{13}C$ and $\delta^{15}N$ values for raw consumer data points and mean source data colored by basin. The shapes represent the source type and the horizontal and vertical lines depict standard deviation.

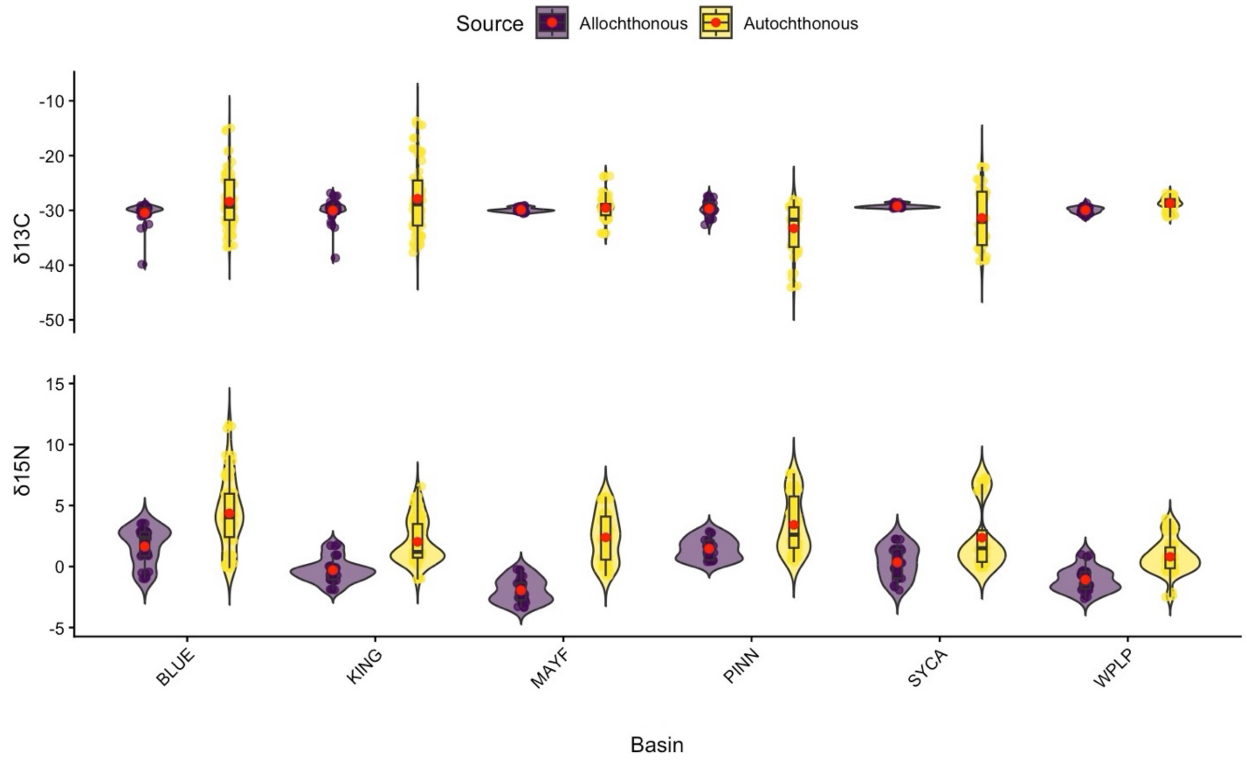


**Figure S4.** Violin plots showing the $\delta^{13}C$ and $\delta^{15}N$ values for raw source data at each basin, colored by source type. Red dots represent the mean value.

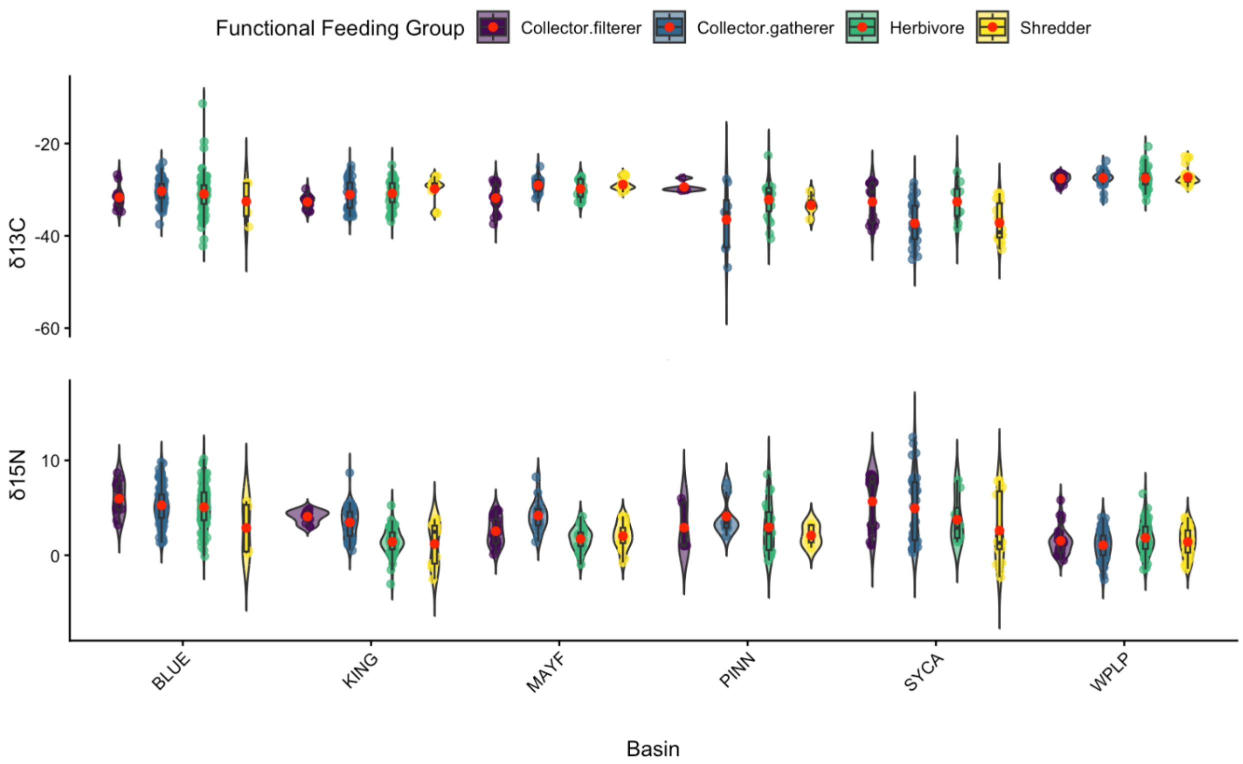


**Figure S5.** Violin plots showing the $\delta^{13}C$ and $\delta^{15}N$ values for raw consumer data at each basin, colored by functional feeding group. Red dots represent the mean value.

(A)

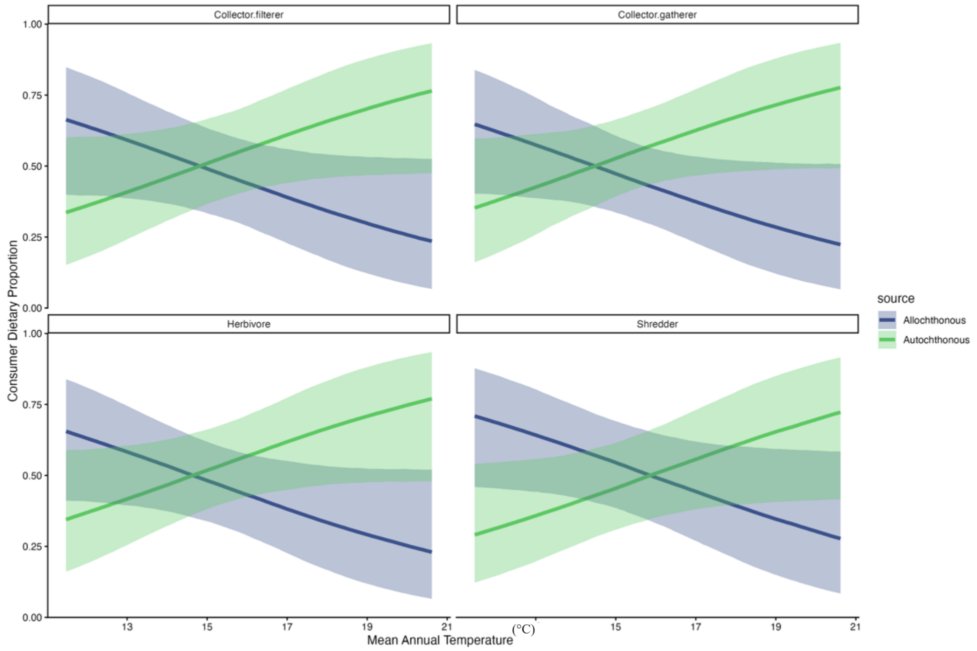


(B)

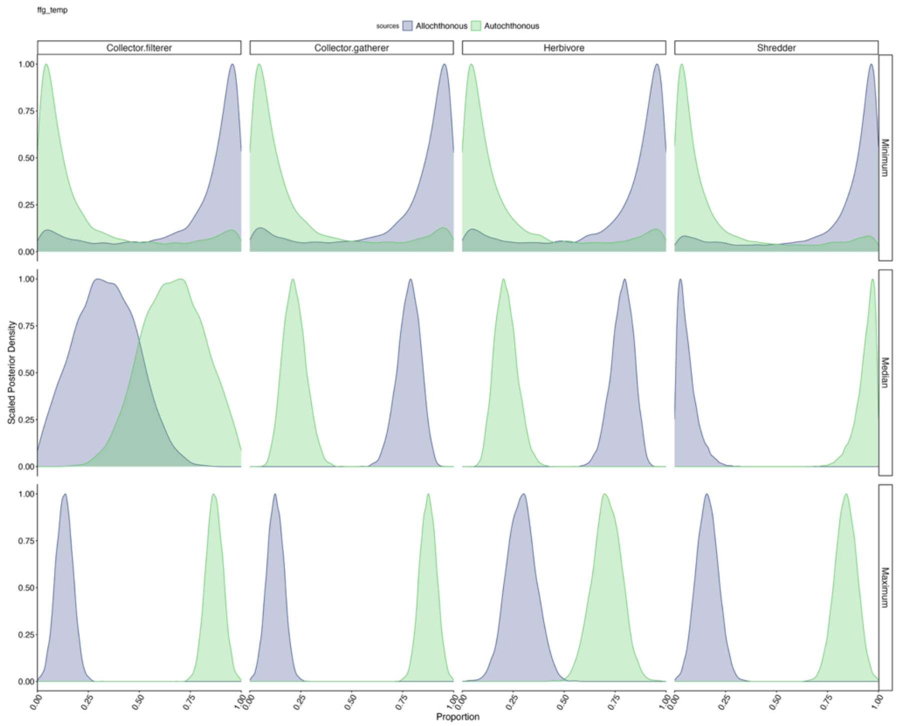


**Figure S6.** (A) Model 7 posterior distribution with 95% credible intervals of consumer dietary proportions of each FFG, plotted against average annual temperature (°C) and colored by source type. (B) Model 7 posterior distributions of dietary proportions for each FFG, at the minimum, median, and maximum average annual temperature (°C), colored by source type.

(A)

(B)

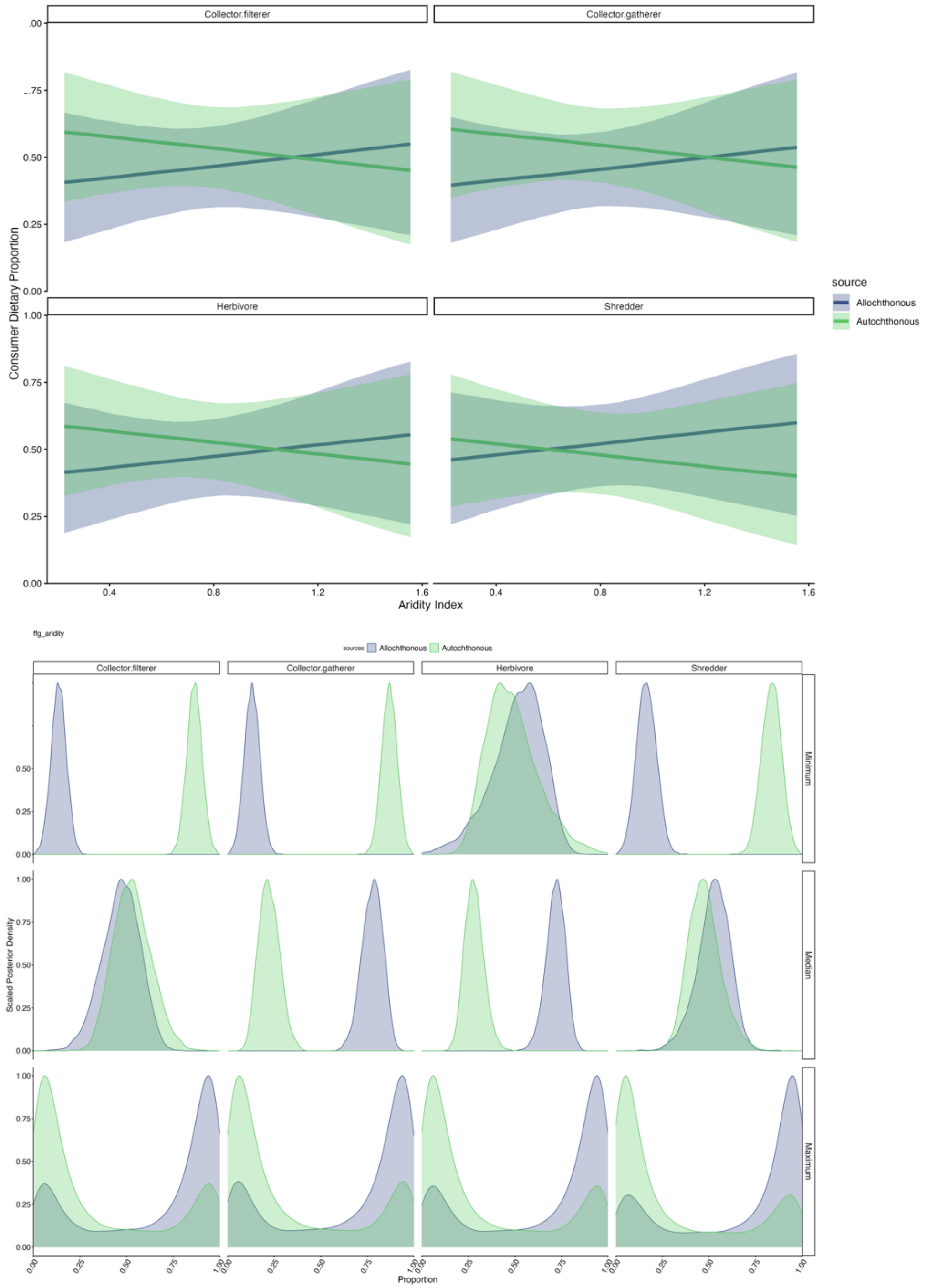


**Figure S7.** (A) Model 9 posterior distribution with 95% credible intervals of consumer dietary proportions of each basin, plotted against aridity index and colored by source type. (B) Model 9 posterior distributions of dietary proportions for each basin, at the minimum, median, and maximum aridity index, colored by source type.

(A)

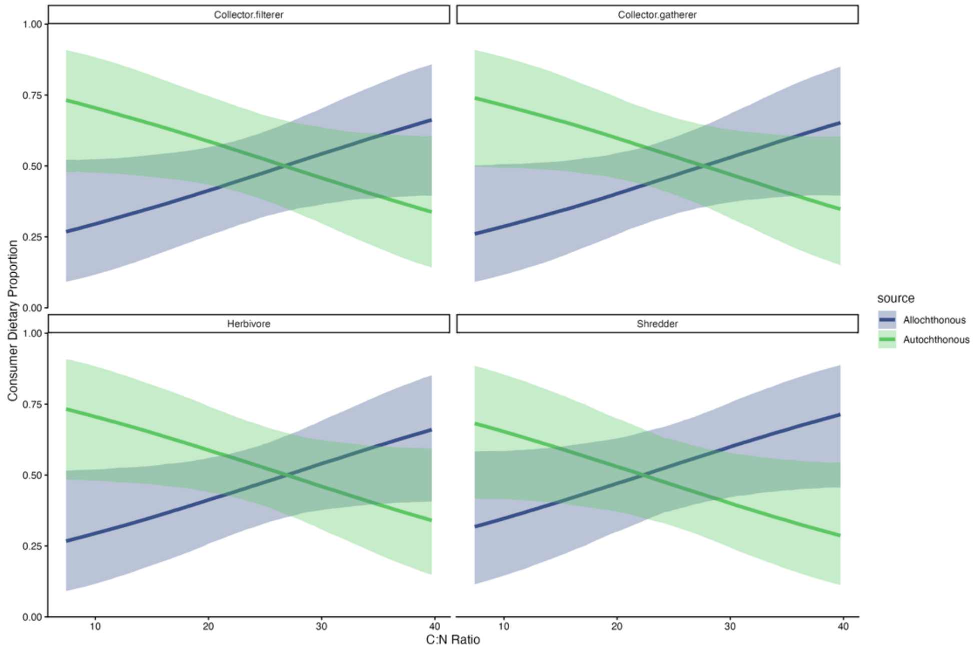


(B)

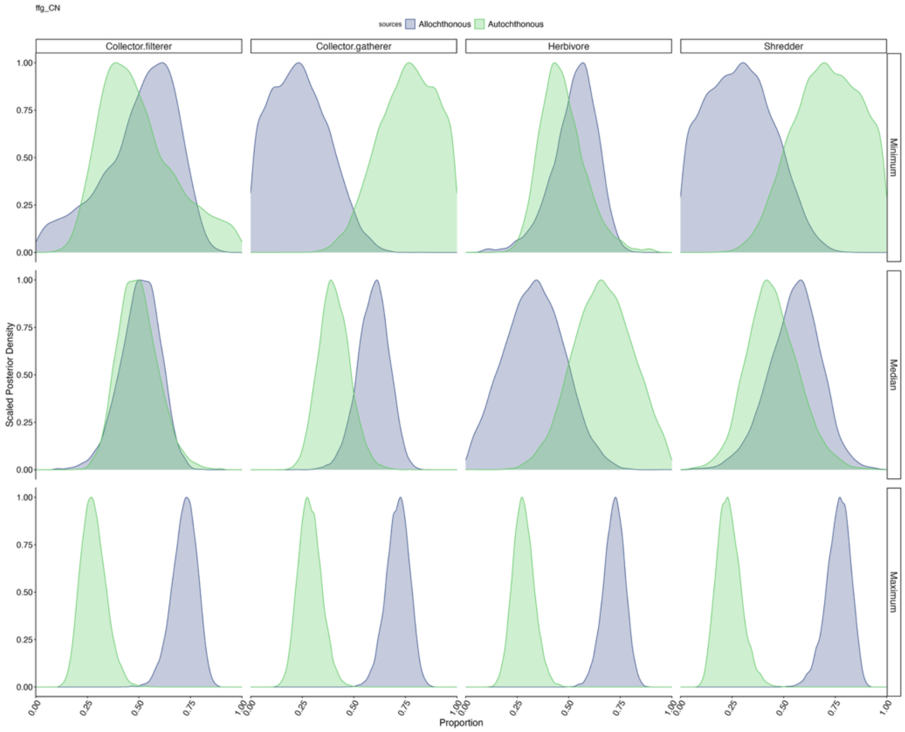


**Figure S8.** (A) Model 10 posterior distribution with 95% credible intervals of consumer dietary proportions of each FFG, plotted against average C:N ratio of allochthonous sources and colored by source type. (B) Model 10 posterior distributions of dietary proportions for each FFG, at the minimum, median, and maximum average C:N ratio of allochthonous sources, colored by source type.

(A)

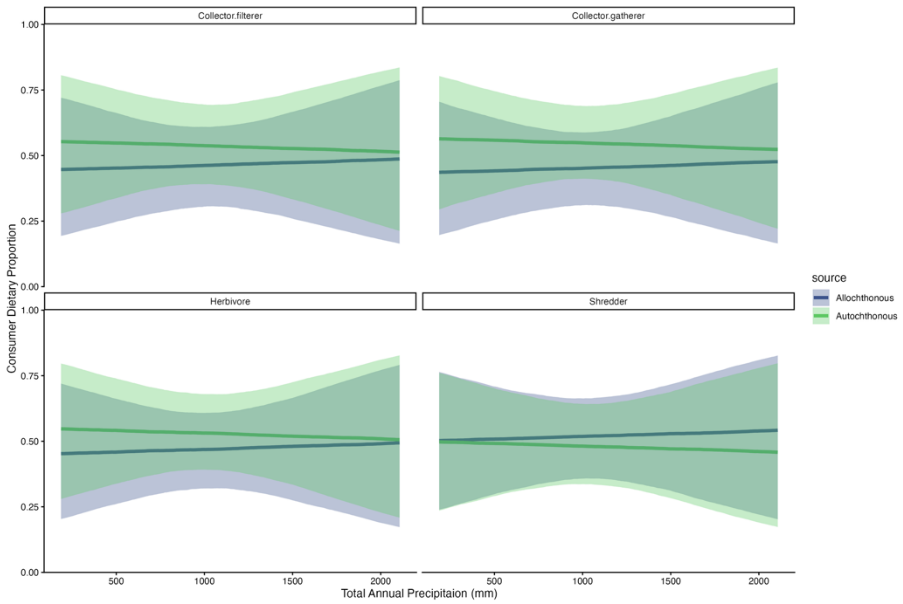


(B)

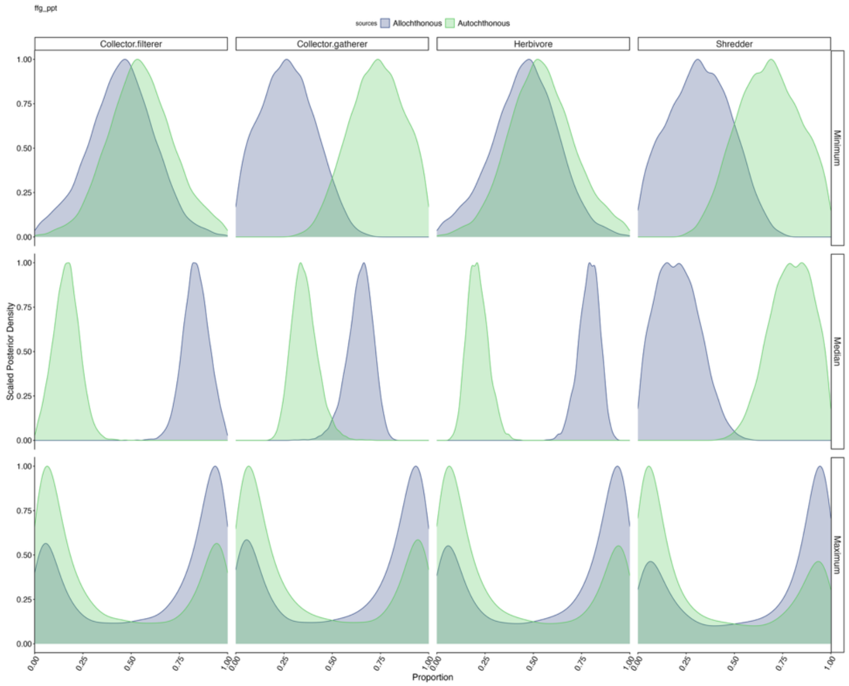


**Figure S9.** (A) Model 8 posterior distribution with 95% credible intervals of consumer dietary proportions of each FFG, plotted against total annual precipitation (mm) and colored by source type. (B) Model 8 posterior distributions of dietary proportions for each FFG, at the minimum, median, and maximum total annual precipitation, colored by source type.

(A)

(B)

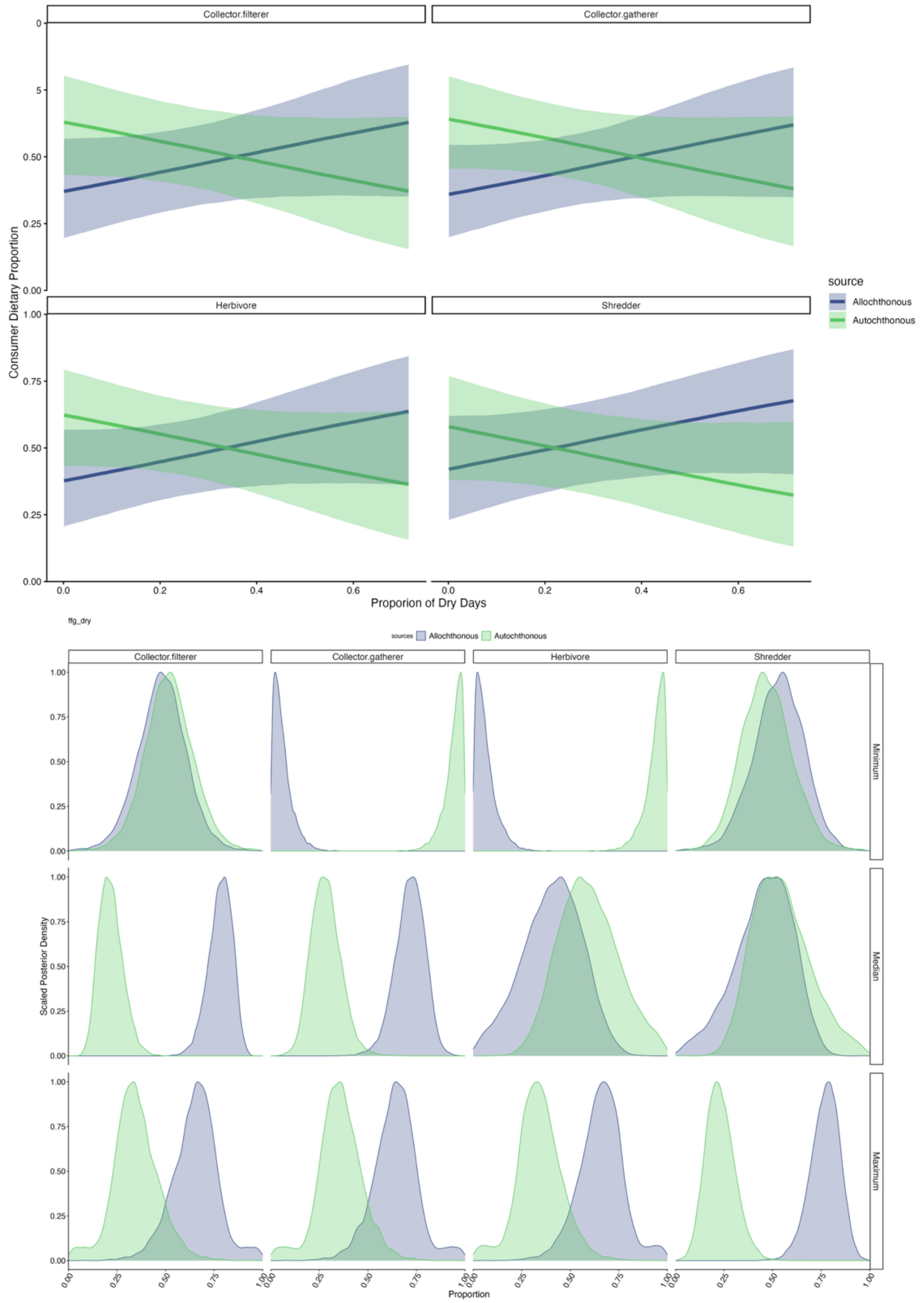


**Figure S10.** (A) Model 5 posterior distribution with 95% credible intervals of consumer dietary proportions of each FFG, plotted against average annual proportion of dry days and colored by source type. (B) Model 5 posterior distributions of dietary proportions for each FFG, at the minimum, median, and maximum proportion of dry days, colored by source type.

(A)

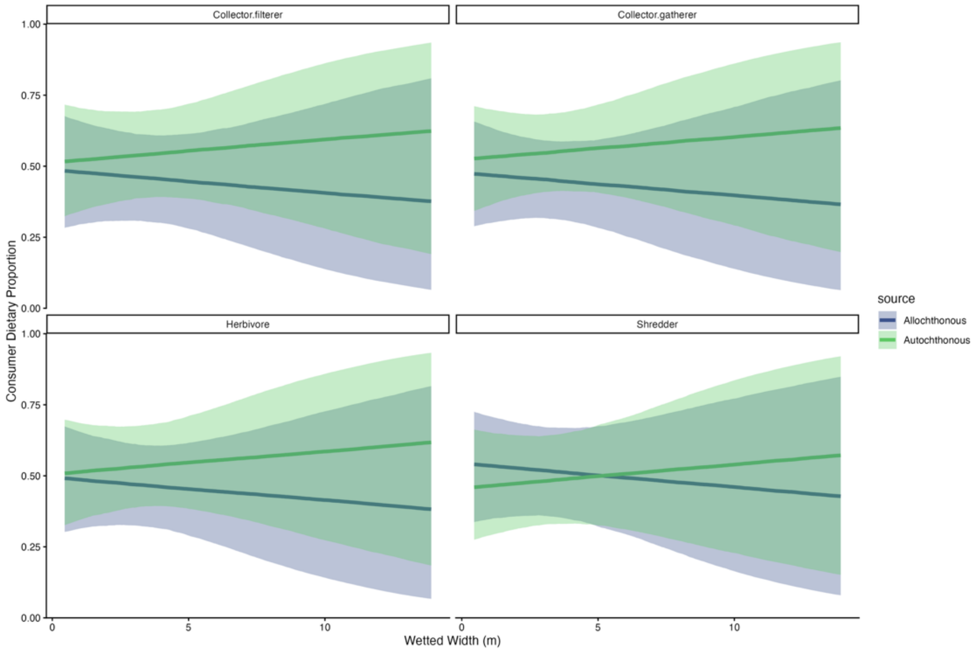


(B)

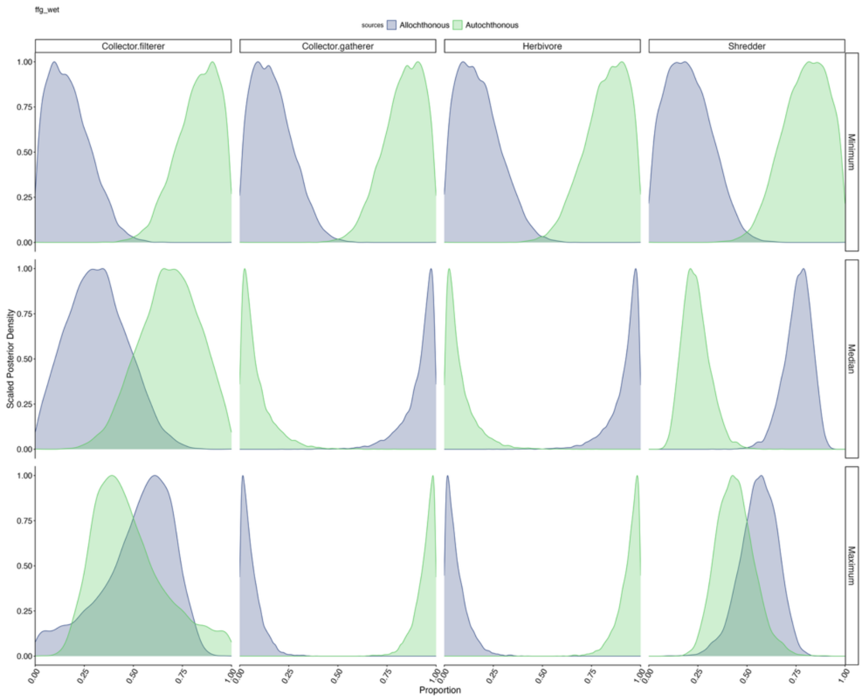


**Figure S11.** (A) Model 4 posterior distribution with 95% credible intervals of consumer dietary proportions of each FFG, plotted against average wetted width (m) and colored by source type. (B) Model 4 posterior distributions of dietary proportions for each FFG, at the minimum, median, and maximum wetted width (m), colored by source type.

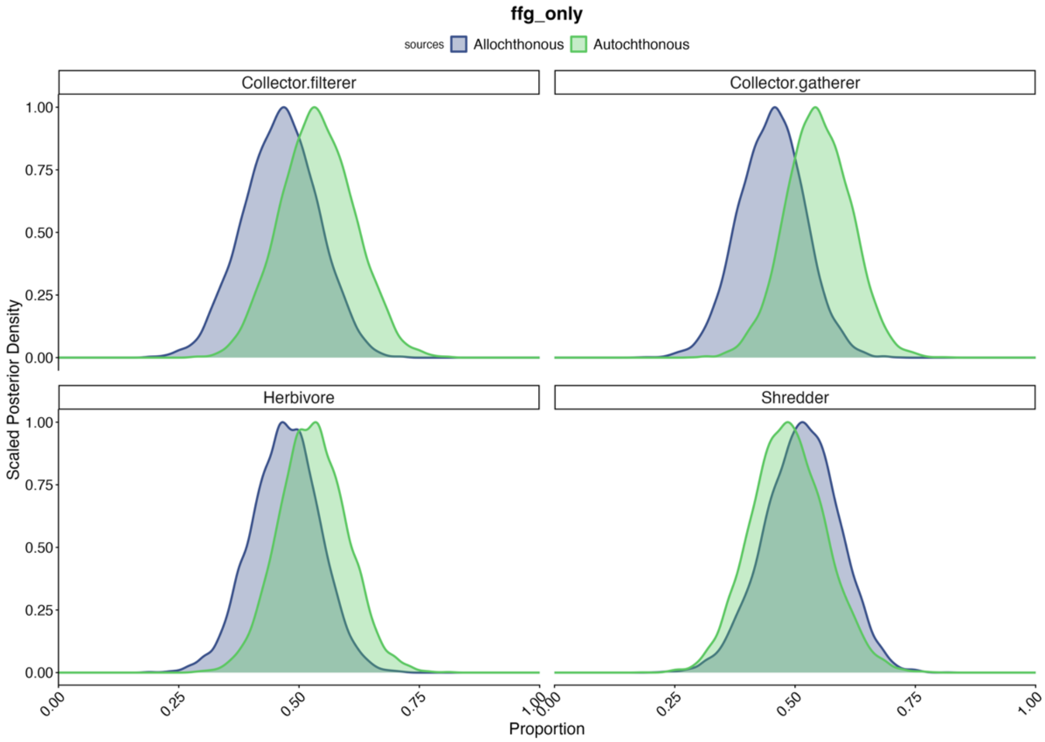


**Figure S12.** Model 2 posterior distributions of dietary proportions of consumers for each FFG.

(A)

(B)

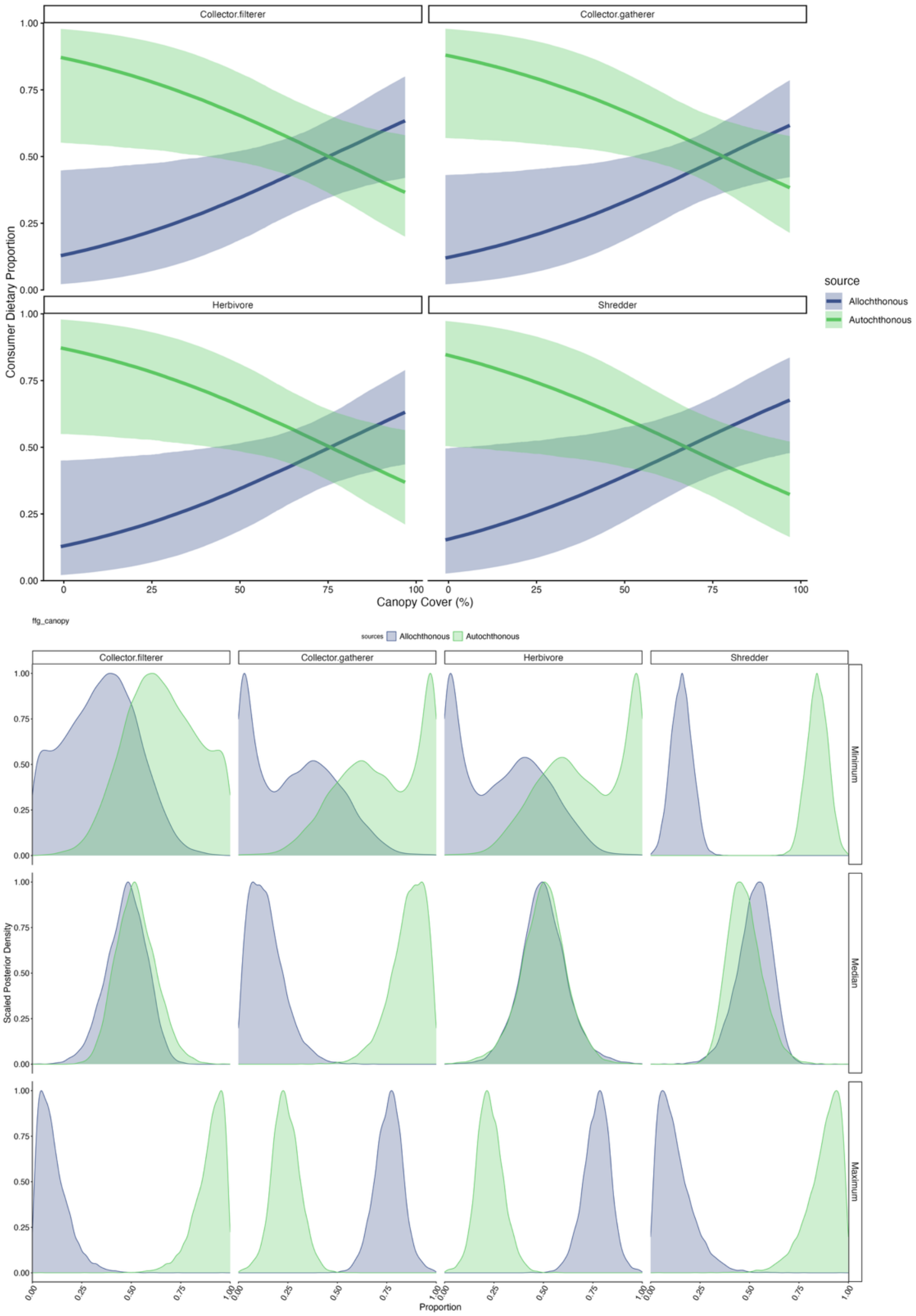


**Figure S13.** (A) Model 3 posterior distribution with 95% credible intervals of consumer dietary proportions of each FFG, plotted against canopy cover (%) and colored by source type. (B) Model 3 posterior distributions of dietary proportions for each FFG, at the minimum, median, and maximum canopy cover (%), colored by source type.

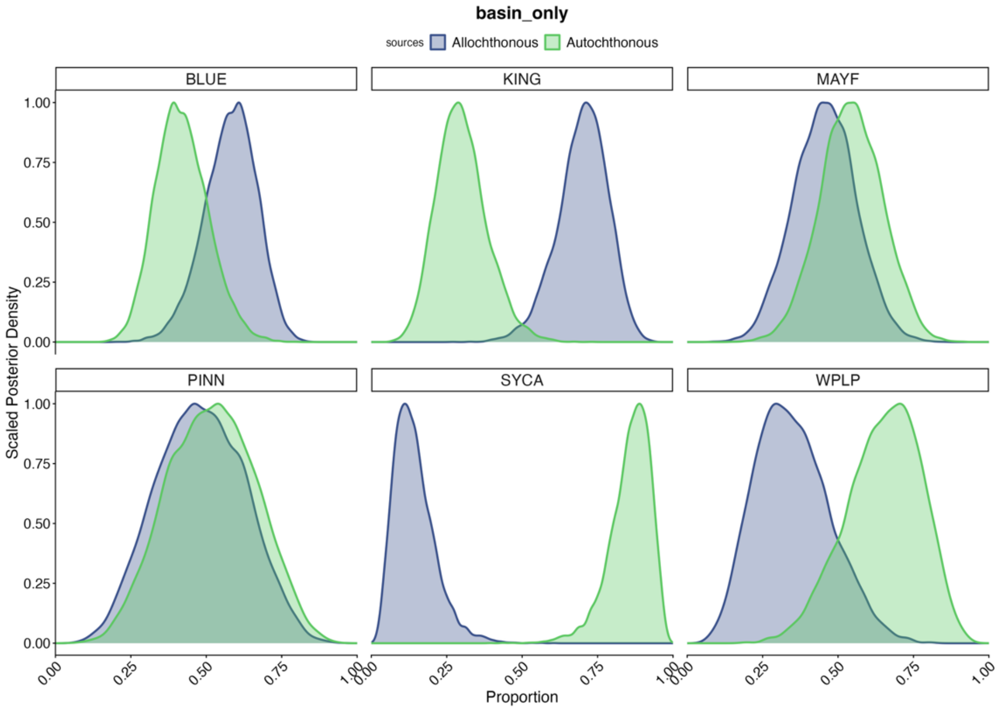


**Figure S14.** Model 1 posterior distributions of dietary proportions of consumers for each basin.

(A)

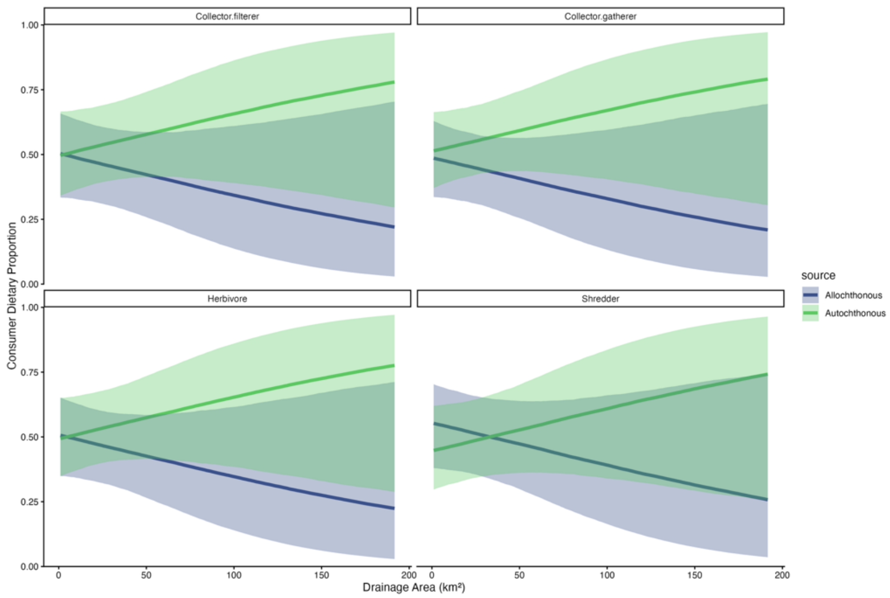


(B)

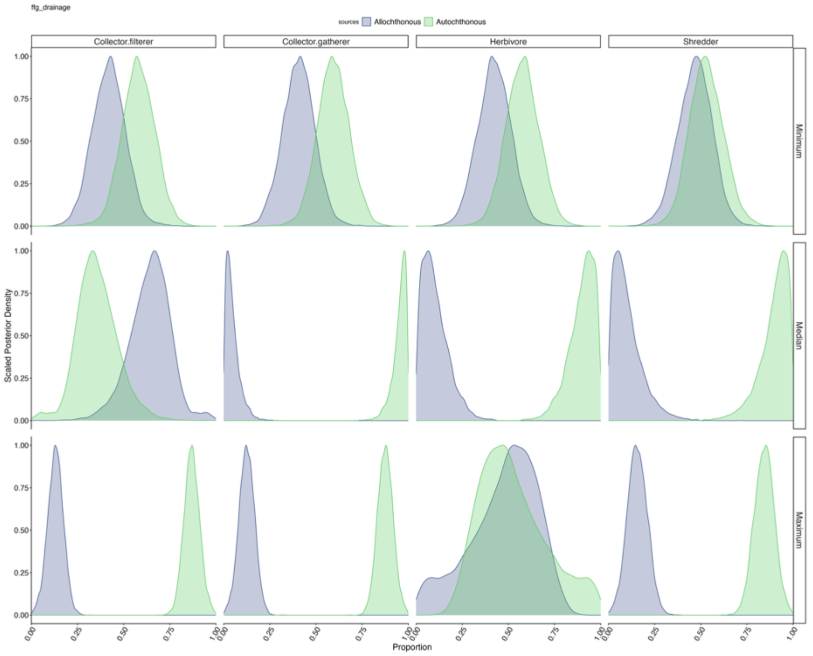


**Figure S15.** (A) Model 6 posterior distribution with 95% credible intervals of consumer dietary proportions of each FFG, plotted against drainage area ($km^2$) and colored by source type. (B) Model 6 posterior distributions of dietary proportions for each FFG, at the minimum, median, and maximum drainage area ($km^2$), colored by source type.
